# Retinal pigment epithelium-derived PD-L1 reprograms microglial cells and protects against retinal degeneration in mouse models of experimental AMD and genetic retinitis pigmentosa

**DOI:** 10.1101/2025.07.26.666839

**Authors:** Zhongyuan Su, Jing Wang, Qinghua Lai, Zhenhang Wang, Pingping Liu, Zhiyan Tang, Hang Yu, Yu Chen, Xiaoyin Ma, Ling Hou

## Abstract

Retinal degeneration is a leading cause of blindness and a significant health burden during aging worldwide. Because of its complex pathogenic mechanisms, however, effective interventions applicable to multiple retinal degenerations remain limited. Excessive microglia-mediated inflammation is a common pathological feature of different retinal degenerative diseases, and inhibiting these overactive inflammatory responses by reprogramming microglia may provide a therapeutic strategy with broad-spectrum potential. Here, we uncover a novel retinal pigment epithelium (RPE) homeostatic signal, PD-L1, that drives the reprogramming of activated microglial toward a retinal-protective phenotype. Using multiple analyses, including gene expression profiling bioinformatics, adeno-associated virus-mediated RPE-specific gene overexpression, pharmacological, and genetic manipulations, we show that RPE cells upregulate the expression of PD-L1 to modulate microglia for protection against retinal degeneration in an experimental dry AMD mouse model. Bioinformatics enrichment analysis of the high-expression gene signatures of subretinal microglia in AMD patients, combined with pharmacological blocking of NF-κB activity or using *Nlrp3* knockout mice, show that PD-L1/PD-1 signaling suppresses NF-κB activation in microglia, thereby inhibiting NLRP3 inflammasome priming and attenuating microglia-mediated excessive inflammation. Furthermore, the overexpression of PD-L1 by the AAV8 virus exerts similar microglial reprogramming and retinal protective effects to alleviate retinal degeneration in a preclinical model of retinitis pigmentosa. These findings reveal that RPE-derived PD-L1 suppresses inflammatory reprogramming of subretinal microglia and mitigates retinal degeneration, highlighting a promising proof of concept for PD-L1 as a potential therapeutic target for the broad treatment of retinal degenerative diseases.

## Introduction

Retinal degeneration is a gradual process characterized by the progressive loss of vision resulting from the failure of photoreceptors and/or of retinal pigment epithelium (RPE). Retinal degeneration is due to inflammation, oxidative stress, environmental factors, and influenced by genetic factors and aging. Retinal degeneration is seen in a group of genetically and clinically heterogeneous disorders (retinal degenerative diseases, RRDs) that include age-related macular degeneration (AMD), inherited retinitis pigmentosa (RP), diabetic retinopathy and glaucoma (Guillonneau et al., 2017; Krady et al., 2005; Tezel, 2022; Wang and Cepko, 2022). The retina, with its photoreceptors, bipolar cells, horizontal cells, ganglion cells as well as glial cells is essential for visual functions but is also highly susceptible to damage due to its limited regenerative capacity. Hence, the slowing of the progress of degeneration and the preservation of endogenous retinal cells is critical (Li et al., 2020., Wang et al., 2019., Wang et al., 2020). In fact, the retina intrinsically employs multiple mechanisms to mitigate structural damage, paramount among which is the reduction of detrimental immune and inflammatory responses (Chen et al., 2019).

Microglia, the resident immune sentinels of the retina, become activated in various RRDs (Karlstetter et al., 2015). Accumulating evidence highlights the central role of excessive neuroinflammation mediated by microglia in disease progression (Guillonneau et al., 2017; Krady et al., 2005; Rao et al., 2003; Tezel et al., 2022; Zabel et al., 2016). Therefore, targeting microglia will offer a promising strategy for preventing or slowing the progression of retinal degeneration. Potential therapeutic strategies include depletion, effector blockage, and microglial reprogramming (Wang and Cepko, 2022), which have shown promising results in the treatment of both experimental autoimmune uveoretinitis (EAU) and inherited retinal degeneration (Okunuki et al., 2019; Zhao et al., 2015; Wang et al., 2020). However, due to the heterogeneity and complexity of its functions, the mechanisms that generate microglial responses during retinal degeneration are not fully understood.

There is increasing evidence to suggest that microglial cells play a crucial role in maintaining retinal synaptic connections (Karlstetter et al., 2015; Wang et al., 2016). Their different subpopulations exhibit distinct microenvironment-dependent protective behaviors (O’Koren et al., 2019) that may be contradictory. For instance, they may exacerbate retinal damage through dysregulated inflammatory responses or inappropriate phagocytosis of viable neurons (Zhao et al., 2015). Notably, the depletion of these cells using CSF1R inhibitors, such as PLX5622, is effective in attenuating the progression of experimental autoimmune uveoretinitis (EAU) (Okunuki et al., 2019). However, in ocular hypertension-induced glaucoma models, microglial depletion by treatment with PLX5622 did not significantly impact retinal ganglion cell degeneration or preservation of visual function (Hilla et al., 2017; Tan et al., 2021) nor did PLX5622 or another CSF1R inhibitor, PLX3397, substantially ameliorated secondary cone degeneration in rd1 mice (Wang et al., 2019; Wang et al., 2020). Notably, microglia-specific galectin-3 ablation led to impaired phagocytosis and subsequent exacerbation of photoreceptor death, RPE damage, and visual impairment (Yu et al., 2024). These occasionally contradictory outcomes arise from the plasticity of microglia, whose morphological and physiological phenotypes are dynamically shaped by the pattern of retinal injury and anatomical localizations, thus highlighting the critical influence of local microenvironmental factors and presenting a fascinating challenge to understanding and treating retinal degeneration.

The remarkable adaptability of microglia has spurred the investigation of innovative reprogramming strategies aimed at enhancing their beneficial neuroprotective potential while suppressing their detrimental effects (Scholz et al., 2015; Wang et al., 2020). Approaches to modulate microglial function by reprogramming molecular targets, such as GLP-1R and TGF-β1 signaling, have shown potential in alleviating retinal degeneration and protecting vision (Hernández et al., 2016; Wang et al., 2020; Wang and Cepko, 2022). Single-cell RNA sequencing has revealed that healthy adult mouse retinas comprise at least two functionally distinct populations of microglia: IL-34-dependent and IL-34-independent (O’Koren et al., 2019), while damaged tissue can develop up to seven distinct subpopulations (Yu et al., 2024). These findings suggest that variations in disease-initiating factors and injury-induced microenvironments in different retinal degenerative diseases may lead to distinct microglial subpopulation profiles, implying that the key signals and regulators required for microglial reprogramming may vary significantly depending on the specific type of retinal degeneration. Therefore, discovering relatively broad-spectrum strategies and reprogramming microglia using regulators and signals that work across retinal degenerative diseases carries tremendous importance.

The RPE is a polarized monolayer of pigment cells that separates the neural retina from Bruch’s membrane and the choroid, playing critical roles during eye development, subretinal homeostasis, and visual function (Strauss 2005). Besides its well-known functions in maintaining retinal health, including clearance of photoreceptor debris, managing the visual cycle, producing neurotrophic factors, and providing antioxidant protection (Strauss 2005; Ma et al., 2019), it is also essential for creating the immune-privilege of the posterior eye (Chen et al., 2019), helps form the blood-retina barrier and restrains T-cell activity (Sugita et al., 2006; Idelson et al., 2018). Dysfunction or loss in RPE cells often leads to retinal degeneration and visual impairment, such as seen in AMD (Fleckenstein et al., 2021). In AMD cases, a specialized group of microglia is recruited to the subretinal area, where they encounter a unique microenvironment rich in photoreceptor debris and injured RPE (Guillonneau et al., 2017). Intriguingly, upon interacting with the injured subretinal microenvironment, these microglia exhibit divergent effects on the RPE. On the one hand, microglia exacerbate inflammatory responses by upregulating RPE chemokine expression through IL-1β signaling, therefore aggravating retinal dystrophies (Natoli et al., 2017). On the other hand, impairment of the phagocytic activity of microglia following galectin-3 deficiency exacerbates RPE damage and retinal degeneration, revealing that microglia play a crucial role in maintaining RPE homeostasis through efficient debris clearance (Yu et al., 2024). These seemingly contradictory findings suggest that subretinal microglia possess both neurotoxic and neuroprotective capacities, with the balance of their functions critically determining their impacts on RPE and photoreceptor viability. Given the central role of the RPE in retinal immune regulation and its anatomical proximity to subretinal microglia, it is critical to understand how these subretinal microglia interact with damaged RPE cells and how microglia play a complicated role in protection from, or enhancement of, retinal degeneration. Therefore, identifying critical homeostatic signals has crucial therapeutic implications.

Signaling by the programmed death-ligand 1 (PD-L1, encoded by CD274) and its receptor PD-1 represents a critical immune checkpoint that maintains immune tolerance and prevents autoimmunity (Gianchecchi et al., 2013; Sun et al., 2018). PD-L1, expressed on antigen-presenting cells and various parenchymal cells, binds to PD-1 on T cells to inhibit their activation through SHP-2-mediated dephosphorylation of key signaling molecules (e.g., ZAP70) (Hui et al., 2017). This immune checkpoint pathway has been implicated in the pathogenesis of various diseases, including neurodegenerative diseases. For instance, in Alzheimer’s disease (AD), PD-L1 is upregulated in reactive astrocytes and may potentially suppress neuroinflammation and Alzheimer’s disease pathology (Kummer et al., 2021). However, the role of the PD-L1/PD-1 checkpoint in microglial inflammatory responses during retinal degenerative diseases, a topic of this study, remains largely unexplored.

In the present study, we show that subretinal microglia have protective effects on NaIO3-induced retinal degeneration. Gene expression profiling using bioinformatics approaches, combined with pharmacological blocking of PD-L1/PD-1 interactions, showed that this protection depends on PD-L1 produced by NaIO3-induced stressed RPE cells. Inhibition of PD-L1/PD-1 signaling accelerated damage to both the RPE and photoreceptors, while specific overexpression of PD-L1 in RPE cells using adeno-associated virus provided retinal protection. Furthermore, our results demonstrate that PD-L1 suppresses NF-κB phosphorylation in microglia, thereby downregulating key components of the NLRP3 inflammasome, including NLRP3, IL-1β, and GSDMD, and ultimately ameliorating retinal degeneration. PD-L1 also exerts similar neuroprotective effects in rd10 mice, a genetic model of primary photoreceptor degeneration. These results reveal a mechanistic basis for PD-L1-driven subretinal microglial cells toward a protective phenotype, suggesting a therapeutic strategy targeting PD-L1 as a potential treatment for diverse retinal degenerations.

## Results

### Subretinal microglia protect RPE and photoreceptors in NaIO3-induced retinal degeneration

To investigate how microglial recruitment after RPE damage affects retinal degeneration, we utilized a NaIO3-induced mouse retinal degeneration model, in which a geographic atrophy-like phenotype (a form of dry AMD) is induced due to primary RPE cells and secondary photoreceptor degeneration (Zhao et al., 2017). On day 1 after treatment of 8-week-old C57BL/6J mice (hereafter referred to as WT mice) with 20 mg/kg NaIO3, mild damage to the RPE and neuroretina was observed in the region near the optic nerve, with no significant damage in other areas. On day 7 after treatment, more breakdowns of RPE monolayer cells (RPE breakdowns) in the area near the optic nerve and the central area were observed, and the outer nuclear layer (ONL) was significantly thinned (Figures 1A, B, and C). Nevertheless, on day 14 after NaIO3-induced damage, RPE breakdowns were only slightly increased compared to day 7; however, statistically significant changes in the thickness of the ONL were observed in a few areas of the superior or inferior regions (Figure 1A, B, and D). The above results indicate that during NaIO3-induced retinal degeneration, cellular damage primarily occurs within the first 7 days, after which the degeneration rate of the RPE and neural retina slows down. These results suggest that after initial damage, the retina may activate self-protective mechanisms.

**Figure 1.**
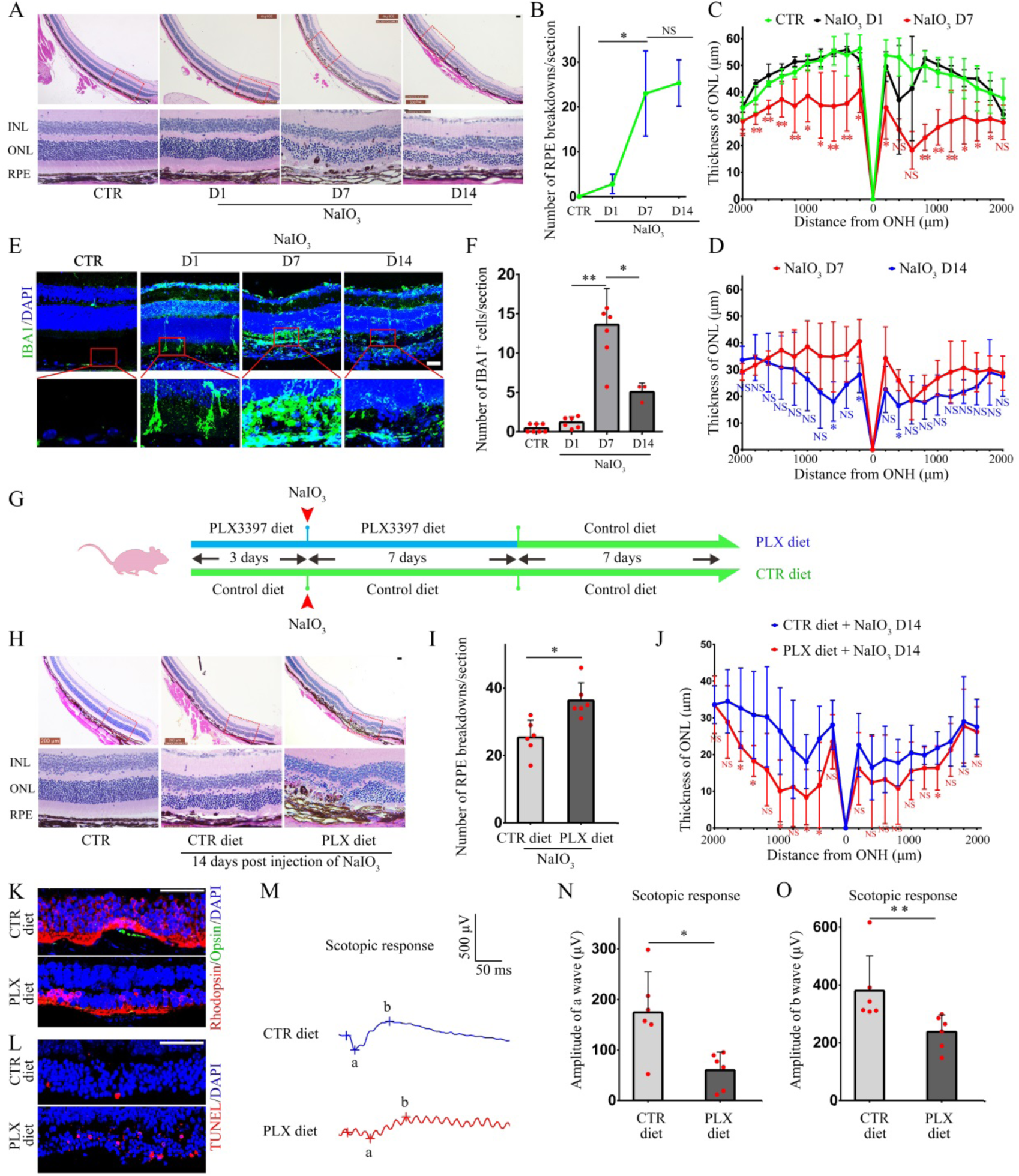
Resident microglia in the subretinal space play a protective role in a NaIO3- induced retina degeneration mouse model. (A) Representative images of H&E staining of retinal structures in WT mice at 1-, 7-, and 14-days post 20 mg/kg NaIO3-induced injury. (B) Quantification of the number of breakdowns of the RPE monolayer cells (RPE breakdowns) based on the results from A, n=6. Note that RPE breakdowns increased rapidly by day 7 compared to controls or day 1, but the increased rate significantly slowed by day 14. (C and D) The curve diagram illustrates the thickness of the ONL based on data from A, n = 6. Note that ONL thickness changes reflected trends of the RPE breakdowns. (E) Representative immunostaining images of IBA1 (green) in WT mice at 1-, 7-, and 14-days after 20 mg/kg NaIO3 injection. Note that the activation of microglia occurs at different time points after the injection of NaIO3. (F) Quantification of the number of subretinal IBA1-positive microglia from E, n = 6. (G) A schematic diagram of microglia depletion using PLX3397 and retinal injury induction, with normally fed mice as controls (CTR). (H) Representative images of H&E staining of retinal structures in WT mice 14 days post 20 mg/kg NaIO3 injection, comparing control and PLX3397-treated groups. (I and J) Quantifications of the number of RPE breakdowns (I) and a curve diagram showing the thickness of ONL (J) based on the results of H, n = 6. (K) Representative immunofluorescence images of Rhodopsin (red) and Opsin (green) double immunostaining showing reduced expression in PLX3397-treated mice. (L) Representative immunofluorescence images of TUNEL assay showing increased apoptosis in the ONL of PLX3397-treated mice. (M) Representative ERG analyses showing reduced scotopic a- and b-wave amplitudes in PLX3397-treated mice. (N and O) Quantifications of scotopic a- and b-wave amplitudes based on the results of M, n = 6. INL, inner nuclear layer; ONH, optic nerve head; ONL, outer nuclear layer; RPE, retinal pigment epithelium. * or ** indicates *P*<0.05 or *P*<0.01. Bar = 50 μm.

Next, we analyzed the distribution of microglia at different stages after NaIO3-induced retinal damage. In the control group, microglia were primarily located in the inner and outer plexiform layers, with almost no microglia present in the subretinal space. However, after one day of treatment with NaIO3, a small number of microglial cells were present in the subretinal space. The presence of microglia at this location peaked on day 7 but was reduced thereafter (Figure 1E, F). This observation was consistent with the alleviation of retinal degeneration over time, suggesting that microglia in the subretinal space play a protective role in RPE cells and photoreceptors. To test whether this may be the case, we depleted microglia using the colony-stimulating factor 1α receptor inhibitor PLX3397 (Figure 1G and Figure S1). Compared with the control chow-fed mice, the number of microglia in the subretinal space was significantly reduced by the PLX3397 treatment. At 14 days after NaIO3 injury, H&E staining showed that the PLX3397-fed group displayed a significantly higher degree of destruction of the RPE (Figure 1H, I) and a significant thinning of the ONL in multiple areas of the superior regions and a few areas of the inferior regions (Figure 1H, J). Moreover, double immunostaining showed reduced expression levels of Rhodopsin, a rod cell marker, and Opsin, a cone cell marker, in the superior area (Figure 1K), indicating that photoreceptor degeneration was more severe after microglia depletion. These results were consistent with the changes in the number of apoptotic cells detected by TUNEL assay. As shown in Figure 1L, although TUNEL-positive cells were present in the ONL of control retinas, their number was much higher in the PLX3397-fed retinas. In addition, electroretinograms (ERGs) showed that while mice treated with NaIO3 for 14 days had a mean scotopic a-wave amplitude of 174.3 μV (range, 52.4-298.1 μV) and a mean b-wave amplitude of 380.7 μV (range, 307.9-616.3 μV), NaIO3-treated/PLX3397-fed mice showed significantly reduced values (mean a-wave amplitude 60 μV, with a range of 11.9-95.3 μV; mean b-wave amplitude 237.8 μV, with a range of 148.1-299.1 μV) (Figure 1 M, N, and O). This finding indicated that the visual function of the retina is more severely impaired after microglial depletion. Taken together, these results suggest that microglia recruited to the subretinal space undergo phenotypic modulation by the local injury microenvironment and acquire protective capabilities, thereby significantly alleviating the structural degeneration and functional decline of RPE cells and photoreceptors in the NaIO3- induced retinal degeneration.

### NaIO_3_-treatment-stressed RPE cells upregulate PD-L1 and blocking its activation exacerbates retinal degeneration

Previous studies have shown that NaIO3-induced retinal damage primarily affects RPE cells, which are the main cellular component of the subretinal injury microenvironment (Moriguchi et al., 2018). To test whether NaIO3-treatment-stressed RPE cells (stressed RPE) may modulate microglia by upregulating specific protective genes, we performed a bioinformatic analysis of previously published gene expression profiles of NaIO3-treated human RPE cells, which mimic phenotypes, including RPE degeneration, observed in AMD (Tang et al., 2021). A comparative analysis revealed that 392 genes were significantly upregulated and 888 genes significantly downregulated in control versus NaIO3-treated groups (Figure 2A). We further analyzed the 392 upregulated genes for the KEGG (Kyoto Encyclopedia of Genes and Genomes) pathway enrichment and found a significant correlation with "PD-L1 expression and the PD-1 checkpoint pathway" (Figure 2B). Key genes in this pathway include *CD274* (encoding PD-L1), *JUN, MAP2K3, JAK2, NFKBIB, TICAM2, FOS, TRAF6, MAPK13*, and *BATF2* (Figure 2C). Given the established immunomodulatory function of the PD-L1/PD-1 checkpoint pathway, we selected PD-L1 for further investigation and confirmed that it was significantly upregulated after RPE cells in cultures were treated with 100 μM NaIO3 for 48 hours (Figure 2D). Western blot analysis further showed that progressive PD-L1 upregulation occurred at both 1 day and 7 days post-NaIO3 injury in RPE cells in mice compared to untreated controls (Figure 2E and F). These results were supported by immunostaining, showing that predominant PD-L1 was expressed in stressed RPE cells and also in a few inner nuclear layer cells 7 days post-treatment with NaIO3 (Figure 2G). This spatial expression pattern suggests that the upregulation of PD-L1 in the subretinal injury microenvironment primarily occurs in RPE cells.

**Figure 2.**
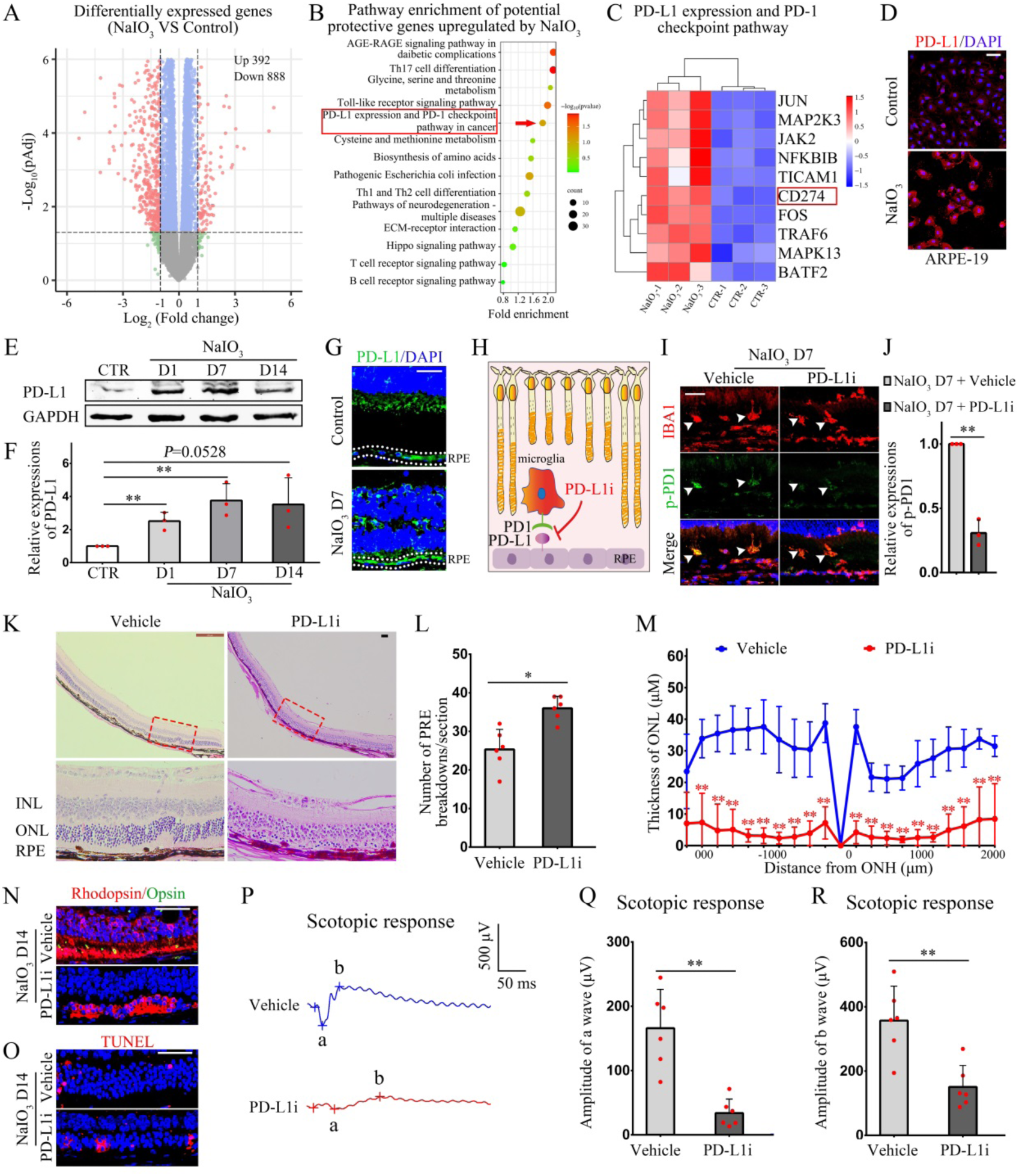
NaIO3-induced RPE stress upregulates PD-L1, and blocking PD-L1 activation promotes retinal degeneration. (A) Analysis of the GSE142591 public dataset from the NCBI Gene Expression Omnibus (GEO) to identify differentially expressed genes, including 392 upregulated and 888 downregulated genes in RPE cells post NaIO3 treatment. (B) KEGG pathway enrichment analysis of upregulated genes, highlighting immune and inflammatory pathways, including PD-L1 expression and the PD-1 immune checkpoint pathway. (C) Heatmap of genes enriched in the PD-L1/PD-1 immune checkpoint pathway. (D) Representative immunostaining images of PD-L1 (red) in ARPE-19 cells with and without NaIO3 treatment, n = 3. (E) Representative western blots measuring levels of PD-L1 in RPE cells in WT mice after NaIO3 injection. (F) The bar graph illustrates the relative expression of PD-L1 based on the results of E, n = 3. (G) Representative immunofluorescence images of PD-L1 (green) showing increased PD-L1 expression in WT RPE after NaIO3 injection. (H) A schematic diagram of PD-L1/PD1 inhibitor 3 (PD-L1i) blocking PD-L1 activation on RPE cells. (I) Representative immunofluorescence images of IBA1 red) and phospho-PD1 (p-PD1, green) showing reduced PD1 phosphorylation in retinal microglia post-PD-L1i treatment, n=3. (J) Quantifications of PD1 phosphorylation levels based on the results of I, n = 3. (K) Representative images of H&E staining showing increased RPE breakdowns and reduced ONL thickness in PD-L1i-treated mice 14 days after NaIO3 injection. (L and M) Quantifications of the number of RPE breakdowns and a curve diagram showing the thickness of the ONL based on the results of K, n = 6. (O) Representative immunofluorescence images of Rhodopsin (red) and Opsin (green) showing reduced expression in PD-L1i-treated mice. (P) Representative immunofluorescence images of the TUNEL assay showing increased apoptosis in PD-L1i-treated mice. (Q) Representative ERG analyses showing reduced scotopic a- and b-wave amplitudes in PD-L1i-treated mice. (R and S) Quantifications of scotopic a- and b-wave amplitudes from the results of Q, n = 6. INL, inner nuclear layer; ONH, optic nerve head; ONL, outer nuclear layer; RPE, retinal pigment epithelium. * or ** indicates *P*<0.05 or *P*<0.01. Bar = 50 μm.

The canonical immunosuppressive mechanism of PD-L1 involves binding to PD-1, which induces PD-1 phosphorylation and activation of downstream signaling. To functionally validate the protective role of RPE-derived PD-L1, we performed subretinal injections of PD-L1/PD1 inhibitor 3 (PD-L1i) in WT mice, effectively blocking PD-L1-mediated PD-1 phosphorylation in microglia (Figure 2H, I, and J). Histological analysis 14 days post-NaIO3 treatment showed that PD-L1i application significantly increased RPE breakdowns (Figure 2K, L) and reduced the thickness of the ONL in both superior and inferior retinal regions (Figure 2K, M). Immunofluorescence results revealed a corresponding decrease in the expression of both Rhodopsin and Opsin (Figure 2N), while TUNEL assays confirmed an increase in apoptotic cells (Figure 2O). In addition, ERG traces of PD-L1i-treated mice were markedly reduced 14 days post-injury: the mean a-wave amplitude was 33.9 μV (range, 13.4-71.4 μV) and the mean b-wave amplitude was 150.6 μV (range, 87.9-268.8 μV), compared with the controls, where the mean scotopic a-wave amplitude was 165.9 μV (range, 82.2-244.5 μV), and the mean b-wave amplitude was 356.9 μV (range, 194.2-509.3 μV) (Figure 2P, Q, and R). Thus, inhibition of PD-L1/PD-1 signaling aggravates NaIO3-induced retinal degeneration. Taken together with our demonstration of injury-induced PD-L1 upregulation in RPE cells, these results support a model in which RPE-derived PD-L1 partially protects RPE cells and attenuates neurodegeneration through PD-L1-mediated modulation of microglial PD-1 phosphorylation.

### PD-L1 overexpression prevents retinal degeneration

The above results, showing that PD-L1/PD-1 inhibition promotes NaIO3-induced retinal degeneration, prompted us to test whether PD-L1 might have a direct protective effect. To induce specific PD-L1 overexpression in RPE cells, we engineered a PD-L1 adeno-associated viral vector, AAV8-pRPE65.-PD-L1-GFP (AAV8-PD-L1), which allows for PD-L1 expression under the control of the RPE65 promoter. AAV8-pRPE65.GFP (AAV8) served as a control vector. When 8-week-old WT mice have subretinally injected with AAV8 or AAV8-PD-L1 virus for three weeks, we confirmed successful infection of RPE cells by GFP expression (Figure S2A). Western blot analysis revealed a significant increase in PD-L1 expression in RPE cells infected with AAV8-PD-L1 compared to the control group infected with AAV8 (Figure S2B, C). Hence, subretinal infection results in a specific overexpression of PD-L1 in RPE cells.

Subsequently, WT mice were infected with AAV8 or AAV8-PD-L1 virus for three weeks, and then subjected to NaIO3-induced injury. Fourteen days post-injury, H&E staining results showed that compared to the control group, RPE and neuroretinal structures in the AAV8-PD-L1-infected group were better preserved, the number of RPE breakdowns was significantly reduced, and the thickness of the ONL was significantly increased (Figure 3A, B, and C). Double immunostaining revealed that the expression levels of Rhodopsin and Opsin proteins were significantly higher than those in the control group, indicating improved preservation of photoreceptors (Figure 3D). TUNEL assays further confirmed a significant reduction in the number of apoptotic cells in the ONL (Figure 3E). Moreover, ERG results showed that in AAV8-infected control mice, the mean amplitude of the scotopic a-wave was 107.9 μV (range, 70.5-166.8 μV), and the mean b-wave amplitude was 319 μV (range, 203.5-371.6 μV). In contrast, scotopic a- and b-wave amplitudes were significantly increased in the AAV8-PD-L1-infected group, with a mean a-wave amplitude of 189.3 μV (range, 148.1-291.8 μV) and a mean b-wave amplitude of 500.7 μV (range, 239.2-667 μV) (Figure 3 F, G, and H). These results indicate that AAV8-mediated overexpression of PD-L1 in RPE cells promotes the preservation of the structure and function of RPE cells and photoreceptors in NaIO3-induced retinal degeneration.

**Figure 3.**
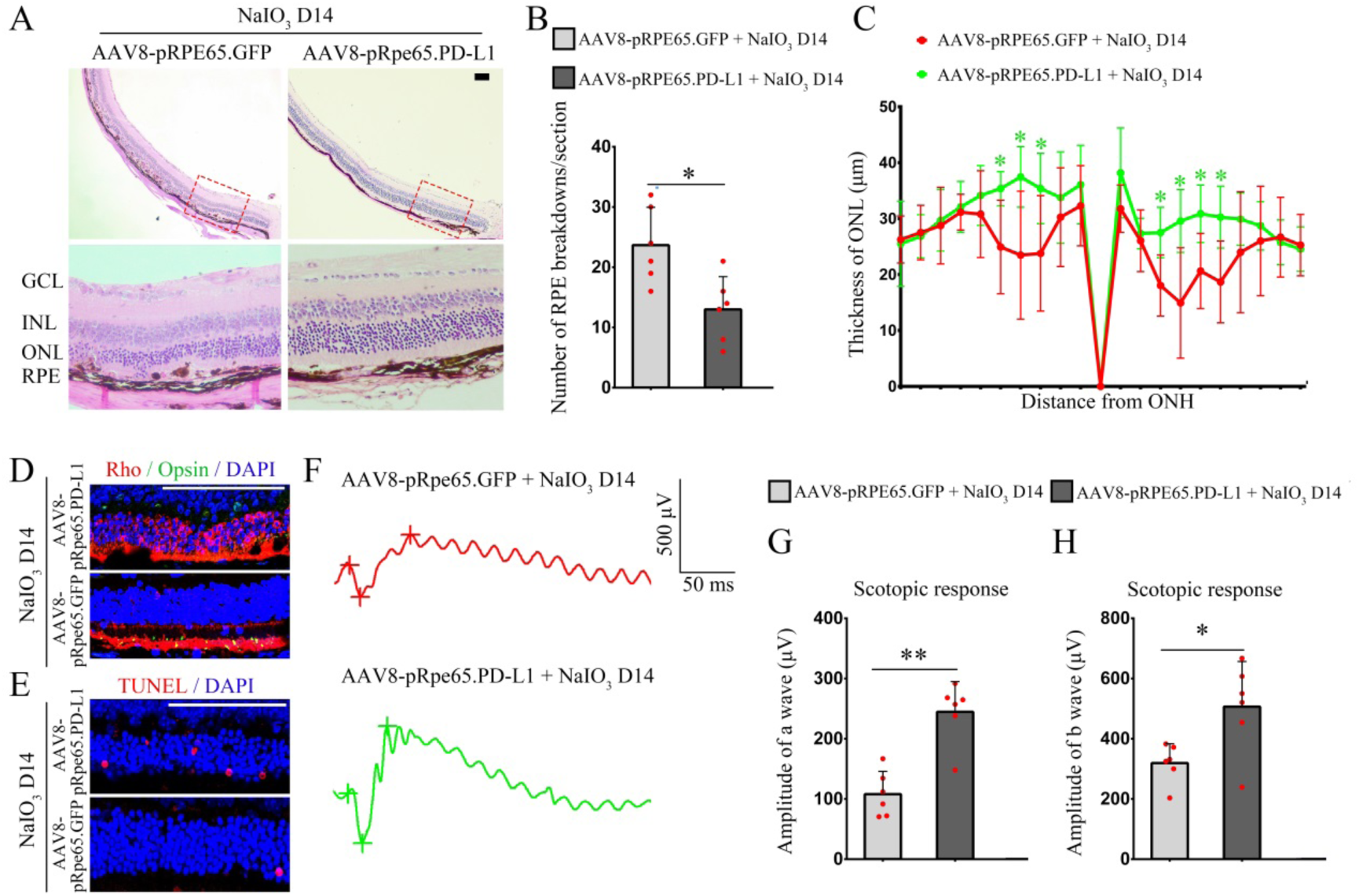
PD-L1 overexpression protects retinas from NaIO3-induced retinal degeneration. (A) Representative images of H&E staining showing better preservation of RPE and retinal structure in AAV8-pRPE65.PD-L1-infected mice 14 days post NaIO3 injury. (B and C) Quantifications of the number of RPE breakdowns and the curve diagram showing the thickness of ONL from A, n = 6. (D) Representative immunofluorescence images of Rhodopsin (red) and Opsin (green) showing increased expression and better preservation of photoreceptors in AAV8-pRPE65.PD-L1-infected mice. (E) Representative immunofluorescence images of the TUNEL assay showing reduced apoptosis in AAV8-pRPE65.PD-L1-infected mice. (F) Representative ERG analyses showing improved scotopic a- and b-wave amplitudes in AAV8-pRPE65.PD-L1-infected mice. (G and H) Quantifications of scotopic a- and b-wave amplitudes from standard response based on the results from F, n = 6. GCL, ganglion cell layer; INL, inner nuclear layer; ONH, optic nerve head; ONL, outer nuclear layer; RPE, retinal pigment epithelium. * or ** indicates *P*<0.05 or *P*<0.01. Bar = 50 μm.

### PD-L1 delays retinal degeneration by inhibiting NF-κB phosphorylation in microglia

The aforementioned results demonstrate that microglia have a protective effect on retinal degeneration after NaIO3-induced retinal injury, and stressed RPE cells upregulate PD-L1 expression, thereby regulating PD-1 phosphorylation in subretinal microglia and delaying retinal degeneration. Given the critical role of the PD-L1/PD-1 pathway in immune regulation, we further investigated how PD-L1 affects the recruitment of subretinal microglia toward a protective phenotype. First, we utilized the human microglial cell line HMC3 to investigate the effect of PD-L1 on the downstream signaling molecules of the PD-L1/PD-1 pathway. After coating culture plates with recombinant human PD-L1 protein, we observed no significant changes in ERK or AKT phosphorylation in HMC3 cells compared to cells cultured in uncoated control dishes. However, as shown in Figures 4A and B, the phosphorylation of the NF-κB p65 subunit was remarkably reduced in the presence of PD-L1. Addition of PD-L1i led to recovery of p65 phosphorylation, while ERK and AKT phosphorylation remained unchanged (Figure 4A, C), suggesting that PD-L1/PD-1 inhibits microglial activation primarily by inhibiting NF-κB p65 phosphorylation.

**Figure 4.**
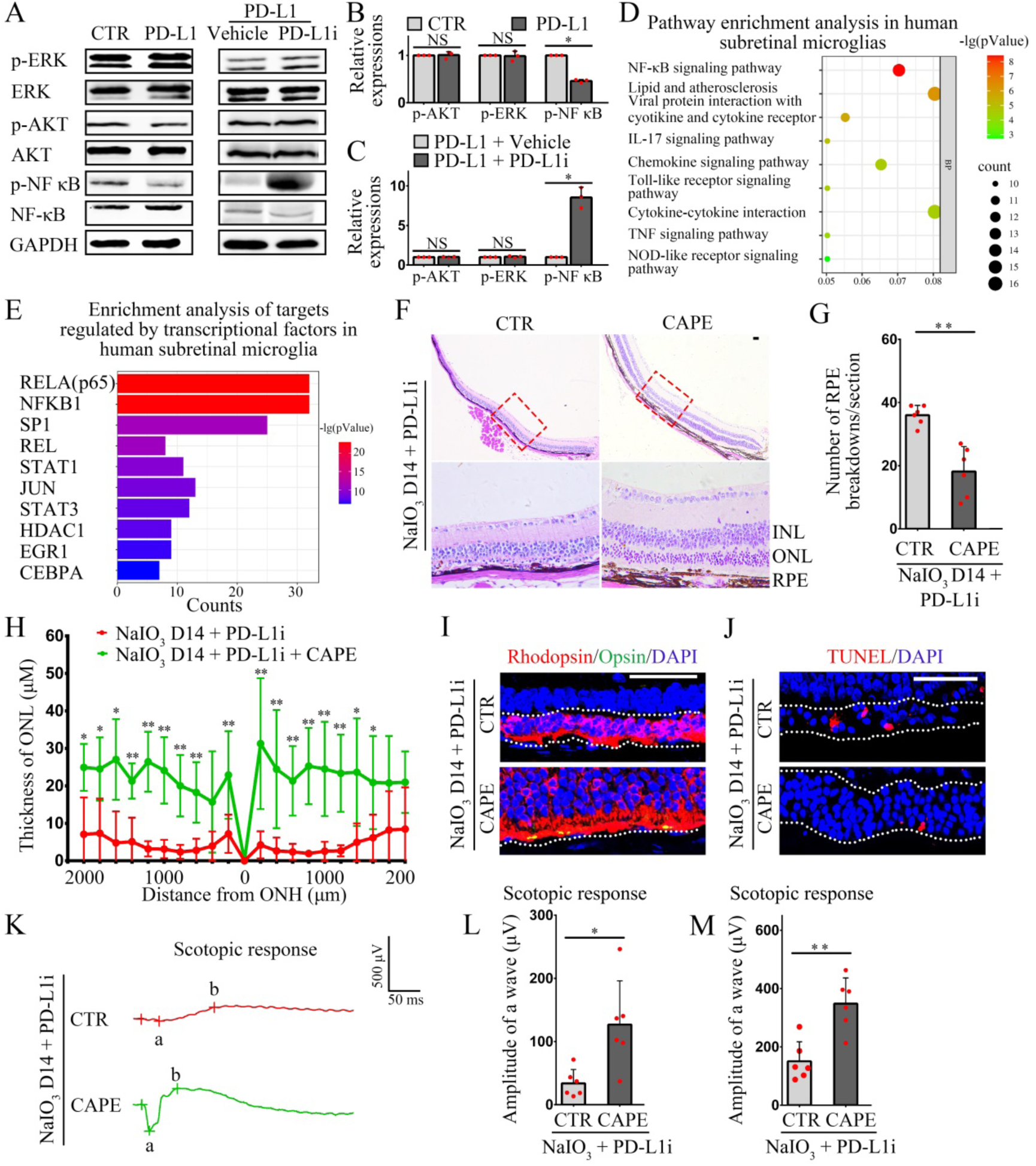
PD-L1 protects retinas by regulating NF-κB phosphorylation in microglia. (A) Western blot analysis of PD-L1’s effects on known PD-1 downstream signaling pathways in human microglial HMC3 cells under indicated conditions. (B, C) Quantifications of phosphorylated AKT, ERK, and NF-κB levels based on results of A, n = 3. (D) The KEGG pathway enrichment analysis of highly expressed genes in subretinal microglia from AMD patients (from Chen et al., 2024). Inflammation-related pathways, notably the NF-κB signaling pathway, were revealed as the top 10 enriched pathways. (E) TRRUST database analysis of transcriptional regulators for the gene set in D revealed significant targeting by NF-κB family members, including RELA (also named as p65), NF-κB1, and REL. (F) Representative images of H&E staining showing better preservation of RPE cells and ONL in PD-L1i + CAPE-treated mice than in PD-L1i alone controls. (G and H) Quantifications of the number of RPE breakdowns and a curve diagram showing the thickness of ONL based on the results of F, n = 6. (I) Representative immunofluorescence images of Rhodopsin (red) and Opsin (green) showing increased expression and better preservation of photoreceptors in PD-L1i + CAPE-treated mice. (J) TUNEL assay showing reduced apoptosis in PD-L1i + CAPE-treated mice. (K) ERG analyses reveal improved scotopic a- and b-wave amplitudes in PD-L1i + CAPE-treated mice, n = 6. Quantifications of a-wave (L) and b-wave (M) amplitudes based on the results from K, n = 6. INL, inner nuclear layer; ONH, optic nerve head; ONL, outer nuclear layer; RPE, retinal pigment epithelium. * or ** indicates *P*<0.05 or *P*<0.01. Bar = 50 μm.

To further assess whether PD-L1 inhibits phosphorylation of NF-κB in microglia, we utilized bioinformatics to analyze the transcriptome of subretinal microglia from AMD patients, based on a previous report (Yu et al., 2024). As shown in Figure 4D, the KEGG pathway enrichment analysis revealed a close association between highly expressed genes in subretinal microglia and inflammatory pathways, with the NF-κB signaling pathway being the most significant enrichment. Intriguingly, the TRRUST database (Transcriptional Regulatory Relationships Unraveled by Sentence-based Text mining) analysis revealed that multiple NF-κB family members, NF-κB1, RELA/p65, and REL, were critical transcriptional regulators of these genes, 32 of which were direct targets of the NF-κB key transcriptional subunit p65 (Figure 4E). These bioinformatic data, combined with our own, suggest that PD-L1/PD-1 pathway inhibits microglia-driven inflammation by suppressing p65 phosphorylation.

Next, we employed caffeic acid phenethyl ester (CAPE), a p65 nuclear translocation inhibitor, for blocking p65 effects (Natarajan et al., 1996). After inducing retinal degeneration with NaIO3, PD-L1i or PD-L1i + CAPE was injected into the subretinal space of mice. The results of nuclear-cytoplasmic fractionation of retinal cell proteins and western blot analysis showed that CAPE significantly inhibited the nuclear translocation of phosphorylated p65 in the PD-L1i + CAPE treatment group compared to that of the PD-L1i only group (Figure S3A, B). The number of RPE breakdowns in the PD-L1i + CAPE group was significantly reduced compared with PD-L1i alone at 14 days of treatment (Figure 4F, G), indicating that inhibiting NF-κB phosphorylation significantly inhibited RPE damage. At the same time, similarly treated mice showed a remarkably greater thickness of the ONL both in superior and inferior regions (Figure 4F, H). In addition, the results of double-immunostaining showed that the expression levels of Rhodopsin and Opsin in the ONL were also significantly higher than those in the control group (Figure 4I), indicating that inhibiting NF-κB phosphorylation preserved more photoreceptors during the degeneration process. TUNEL assay results showed that the number of apoptotic photoreceptor cells was decreased (Figure 4J). These results confirmed that the inhibition of the NF-κB pathway after PD-L1 blockade can significantly attenuate the RPE cell damage and photoreceptor degeneration. In addition, ERG analysis results showed that in the control treatment group, the mean amplitude of scotopic a-wave was 33.9 μV (range, 13.4-71.4 μV), and the mean amplitude of scotopic b-wave was 150.6 μV (range, 87.9-268.8 μV). In contrast, in the PD-L1i + CAPE treatment group, the scotopic a- and b-wave amplitudes were significantly increased, with the mean a-wave amplitude being 126.8 μV (range, 37-246.3 μV), and the mean b-wave amplitude being 348.2 μV (range, 213.2-463.1 μV) (Figure 4K, L, M). Taken together, the above results indicate that stressed RPE cells upregulate PD-L1 to inhibit NF-κB pathway activation in microglia, thereby slowing the progression of retinal degeneration.

### PD-L1 suppresses the expression and activation of NLRP3 inflammasome-associated genes in microglia

Given that microglia have functional plasticity in mediating both pro-inflammatory and anti-inflammatory effects, and that NF-κB is a well-established key regulator of immune responses, we investigated whether PD-L1 may suppress excessive microglial inflammatory responses by inhibiting NF-κB activity, thereby shifting microglia toward a protective phenotype. To do this, we performed a cross-analysis combining the following data: (1) genes reported to be highly expressed in subretinal microglia of AMD patients (Yu et al., 2024), (2) NF-κB-regulated target genes identified by TRRUST analysis, and (3) previously reported NF-κB target genes (Liu et al., 2017). These analyses led to the identification of at least 33 potential NF-κB-regulated target genes that were highly expressed in subretinal microglia, including *NLRP3*, *IL1B*, *BCL2A1*, *IL1A*, *CXCL8*, *CCL2*, *TNF*, and *ICAM1* (Figure 5A). Notably, NLRP3 and IL1B are critical factors involved in NLRP3 inflammasome-mediated inflammatory responses. Moreover, previous studies have shown that the NLRP3 inflammasome plays a pivotal role in driving excessive microglial inflammation (Swanson et al., 2019) and contributes to microglia-mediated neuronal death in Alzheimer’s disease, retinal ganglion cell degeneration, and other neurodegenerative conditions (Puyang et al., 2016; Ising et al., 2019; Wan et al., 2020). Hence, we focused on the role of PD-L1 in regulating the NLRP3 inflammasome-mediated excessive inflammatory responses in microglia. Using the solvent-treated groups as a control, we injected PD-L1i into WT mice and subsequently induced retinal degeneration with NaIO3. Next, we used ionized calcium-binding adapter molecule 1 (IBA1) to label microglia and examined the expression of NLRP3 inflammasome-associated proteins. Seven days post-NaIO3 injection, the control groups showed microglial accumulation in the subretinal space, with clear expression of NLRP3, IL-1β, and GSDMD, indicating that the NLRP3 inflammasome was activated. In addition, dot-like GSDMD aggregations were observed in some microglia (Figure 5B), suggesting that subretinal microglia were undergoing pyroptosis. Upon PD-L1i treatment, the expression levels of NLRP3, IL-1β, and GSDMD in microglia were significantly enhanced (Figure 5B, C), indicating that the inhibition of PD-L1/PD1 signaling further promoted the activation of the NLRP3 inflammasome. This may lead to increased microglial pyroptosis and the release of pro-inflammatory cytokine factors, such as IL-1β, thereby potentially recruiting additional microglia to promote inflammatory responses.

**Figure 5.**
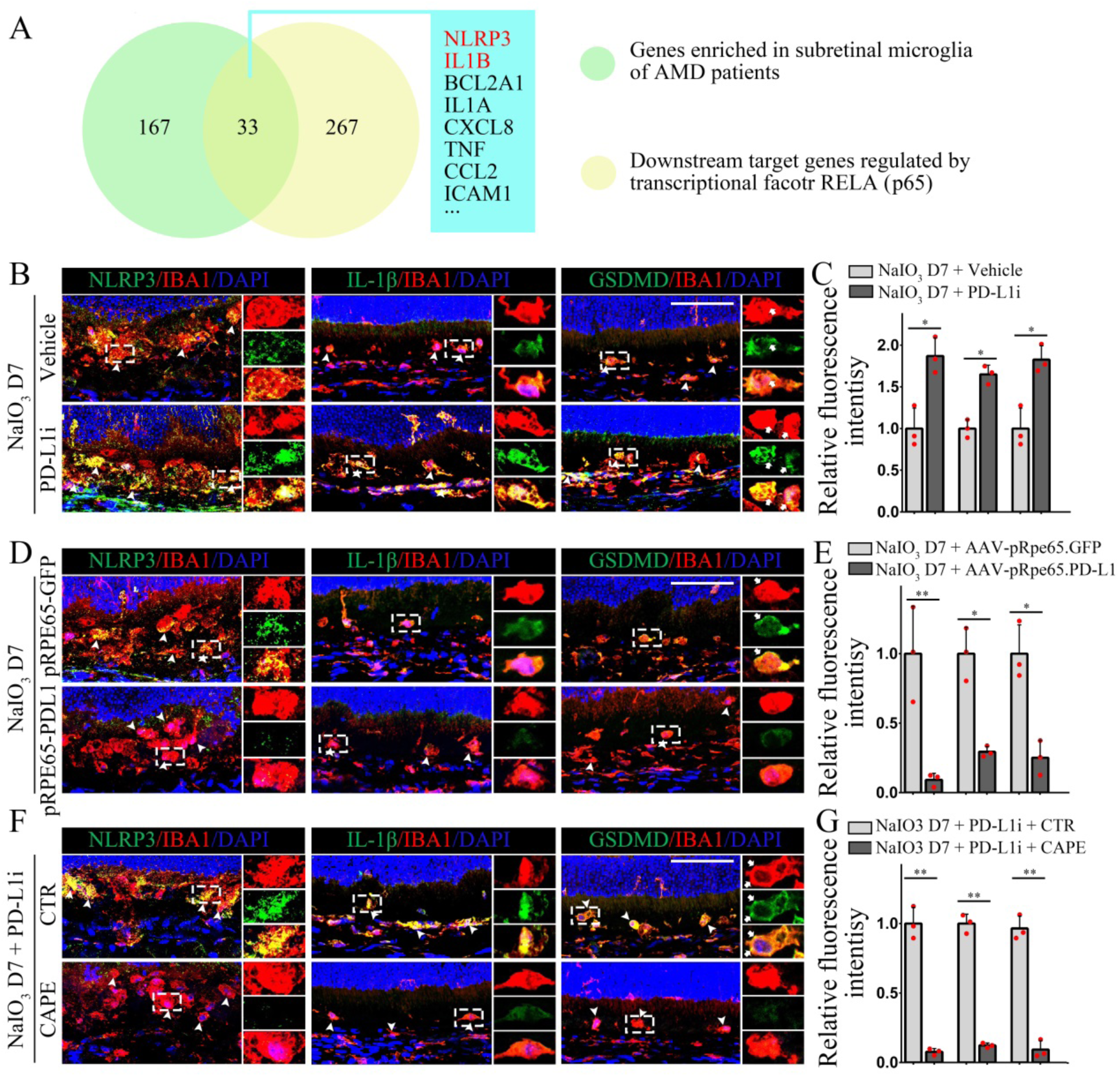
PD-L1 regulates NLRP3 inflammasome-related gene expression and activation in microglia. (A) Bioinformatics analysis of potential NF-κB target genes modulated by PD-L1 in subretinal microglia. Integration of 299 putative p65 targets (from TRRUST database and a literature review of NF-κB signaling) with highly expressed genes in microglia of AMD patients (Chen et al., 2024) identified 33 potential NF-κB-regulated genes, including *NLRP3*, *IL1B*, *BCL2A1*, *IL1A*, *CXCL8*, *CCL2*, *TNF*, and *ICAM1*. (B) Representative immunofluorescence images of IBA1 (red) and NLRP3 (green), IBA1 (red) and IL-1β (green), and IBA1 (red) and GSDMD (green) in microglia after NaIO3 injection and further enhanced by PD-L1i treatment. (C) Quantification of NLRP3, IL-1β, and GSDMD expressions based on the results of B, n = 3. (D) WT mice received subretinal injections of adeno-associated virus AAV8-pRPE65-PD-L1 or control virus AAV8-pRPE65-GFP. Three weeks post-injection, retinal degeneration was induced by NaIO3. Representative immunofluorescence images of IBA1 (red) and NLRP3 (green), IBA1 (red) and IL-1β (green), and IBA1 (red) and GSDMD (green) in microglia at the 7^th^ day post NaIO3-induced injury show reduced NLRP3, IL-1β, and GSDMD expression in AAV8-pRPE65-PD-L1-infected mice. (E) Quantification of relative fluorescence intensities of NLRP3, IL-1β, and GSDMD based on the results of D, n = 3. (F) Representative immunofluorescence images of IBA1 (red) and NLRP3 (green), IBA1 (red) and IL-1β (green), and IBA1 (red) and GSDMD (green) in microglia upon treatment of PD-L1i and CAPE 7 days post NaIO3-induced injury. (G) Quantifications of relative fluorescence intensities of NLRP3, IL-1β, and GSDMD based on the results of F, n = 3. * or ** indicates *P*<0.05 or *P*<0.01. Bar = 50 μm.

Furthermore, we explored the potential of PD-L1 overexpression to alleviate excessive inflammatory responses. As shown in Figures 5D and E, compared with AAV8 controls in WT mice, the expression levels of NLRP3, IL-1β, and GSDMD in microglia were significantly reduced with AAV8-PD-L1 at day 7 post-NaIO3 treatment. These data indicate that the overexpression of PD-L1 in RPE cells could effectively suppress the activation of the NLRP3 inflammasome in subretinal microglia, thereby potentially alleviating excessive inflammatory responses.

Building on our understanding of NF-κB as a critical transcriptional regulator of NLRP3 inflammasome genes and its role in mediating the effects of PD-L1 on retinal degeneration, we utilized CAPE to examine NaIO3-induced retinal injury, thereby combining PD-L1 inhibition with NF-κB suppression. The results, as shown in Figures 5F and G, revealed a significant reduction in the expression levels of NLRP3, IL-1β, and GSDMD in microglia in the PD-L1i + CAPE group compared to the control group using PD-L1i alone. These data suggest that PD-L1/PD1 signaling protects against retinal degeneration by inhibiting the expression of NLRP3 inflammasome-associated genes, primarily through the inhibition of NF-κB in microglia.

### NLRP3 deletion ameliorates retinal degeneration after inhibiting PD-L1 function

In this study, we aimed to investigate the role of NLRP3 in retinal degeneration and its potential interaction with PD-L1. To provide strong genetic evidence, we employed 8-week-old *Nlrp3* gene knockout mice (hereafter referred to as *Nlrp3-/-*) and used age-matched WT mice as controls. All mice were treated with the PD-L1i followed by NaIO3-induced retinal degeneration. Seven days after treatment, we conducted IBA1/NLRP3 co-immunostaining to assess the expression of NLRP3 in subretinal microglia. These results revealed robust NLRP3 expression in subretinal microglia of WT retinas, while NLRP3 protein was completely undetectable in microglia of *Nlrp3-/-* retinas (Figure S4).

To investigate the mechanism of NLRP3 inflammasome-mediated PD-L1 regulation in microglia during retinal degeneration, the mice were treated using the aforementioned method, and samples were collected on day 14. As shown in Figures 6A and B, under the condition of PD-L1 inhibition leading to exacerbated retinal degeneration, H&E staining of retinas of *Nlrp3-/-* mice showed a significant reduction in the number of RPE breakdowns compared with the control group, indicating that the NLRP3 deletion significantly alleviated RPE damage. At the same time, ONL in both superior and inferior retinal regions in similarly treated *Nlrp3-/-* mice was significantly thicker than that in the controls (Figure 6A, C). The results of double immunohistochemical staining revealed that expression levels of Rhodopsin and Opsin in the ONL of *Nlrp3-/-* mice were significantly higher than those in the control group (Figure 6D), and the apoptosis rate was significantly reduced (Figure 6E). These results suggest that NLRP3 deletion can prevent retinal degeneration induced by the loss of PD-L1 function. Additionally, ERG analysis results showed that NLRP3 deletion can improve visual function. As shown in Figures 6F, G, and H, in the control groups, the mean scotopic a-wave amplitude was 33.9 μV (range, 13.4-71.4 μV) and the mean scotopic b-wave amplitude was 150.6 μV (range, 87.9-268.8 μV). In contrast, in the *Nlrp3-/-* group, both scotopic a- and b-wave amplitudes were significantly increased, with the mean a-wave amplitude being 155.2 μV (range, 59.1-275.5 μV) and the mean b-wave amplitude being 433.6 μV (range, 253.2-588.4 μV). Collectively, these results suggest that inhibiting the NLRP3 inflammasome can mitigate the excessive microglial inflammatory responses triggered by PD-L1 inhibition, thereby alleviating the associated retinal degeneration.

**Figure 6.**
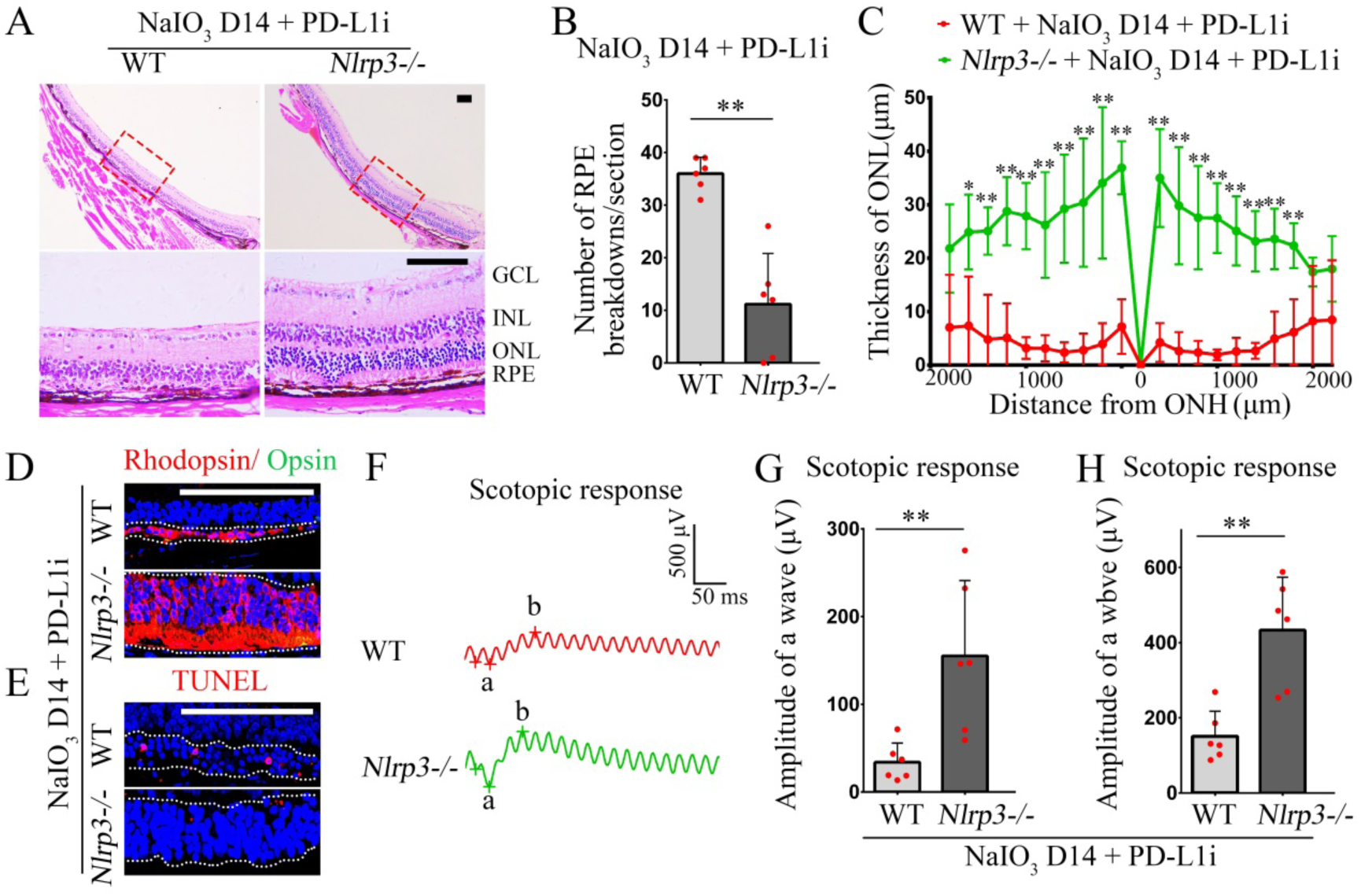
Deletion of NLRP3 in *Nlrp3-/-* mice ameliorates excessive microglial inflammation and retinal degeneration caused by PD-L1 inhibition. All mice were treated with PD-L1i during NaIO3-induced retinal degeneration. (A) Representative images of H&E staining show better preservation of the RPE and ONL in *Nlrp3-/-* mice. (B and C) Quantifications of the number of RPE breakdowns and the curve diagram showing the thickness of the ONL, as determined from the results in A. N = 6. (D) Representative immunofluorescence images of Rhodopsin (red) and Opsin (green) immunofluorescence showing increased expression and better preservation of photoreceptors in *Nlrp3-/-* mice. (E) Representative immunofluorescence images of TUNEL assay showing reduced apoptosis in *Nlrp3-/-* mice. (F) Representative ERG analyses showing improved scotopic a- and b-wave amplitudes in *Nlrp3-/-* mice. (G and H) Quantifications of a- and b-wave amplitudes based on the results of K, n = 6. GCL, ganglion cell layer; INL, inner nuclear layer; ONH, optic nerve head; ONL, outer nuclear layer; RPE, retinal pigment epithelium. * or ** indicates *P*<0.05 or *P*<0.01. Bar = 50 μm.

### PD-L1 overexpression in the retinas of rd10 mice inhibits microglial NLRP3 inflammasome activation, thereby mitigating retinal degeneration

As the above results demonstrated, stressed RPE-derived PD-L1 upregulation modulates subretinal microglial behavior and function, conferring neuroprotective effects in experimental AMD. To further investigate whether overexpression of PD-L1 can protect against genetic retinal degeneration, we subsequently selected the rd10 mouse model. The rd10 mouse model is a widely used model for studying inherited retinal degenerative diseases due to a disease progression that closely resembles human retinitis pigmentosa (RP) (Cronin et al., 2012). Rd10 mice carry a missense mutation in the *Pde6b* gene, leading to rod photoreceptor degeneration in the ONL beginning at postnatal day 18, followed by secondary cone degeneration, ultimately resulting in only 1-2 remaining cell layers in the ONL and near-complete loss of visual function. However, the relatively rapid disease progression made these mice a less straightforward model for AAV8-mediated PD-L1 overexpression. Hence, to delay retinal degeneration, we maintained rd10 mice under darkrearing conditions as previously described (Cronin et al., 2012). To achieve rapid and high expression of PD-L1 in retinal cells, we constructed an AAV8-pCMV.PD-L1-FLAG virus in which PD-L1 expression is under control of the CMV promoter (hereafter referred to as AAV8-pCMV.PD-L1). Following subretinal injection of either AAV8-pCMV.PD-L1 or control virus AAV8-pCMV.FLAG, immunostaining of FLAG-tagged proteins in rd10 mice at postnatal day 14 was examined and confirmed effective infection of photoreceptor cells by both viruses (Figure S5A). These results were supported by Western blot analysis, which confirmed that PD-L1 expression was significantly increased in retinas, primarily in photoreceptor cells, following AAV8-pCMV administration.PD-L1 virus infection compared to the controls (Figure S5B, C).

Next, we employed the above similar treatment protocols to analyze the expression of NLRP3 inflammasome-related proteins in microglia, labeled with IBA1, at 14 days post-AAV infection. As shown in Figures 7A, C, and E, microglia in the ONL of control-infected retinas exhibited explicit expressions of NLRP3, IL-1β, and GSDMD in microglia, whereas the AAV8-pCMV.PD-L1 group showed a significant reduction in the expression of those proteins (Figure 7A, B, C, D, E, and F). This successful reduction in inflammasome-related gene activation in microglia in rd10 mice confirms the potential of PD-L1 overexpression to inhibit inflammasome activation.

**Figure 7.**
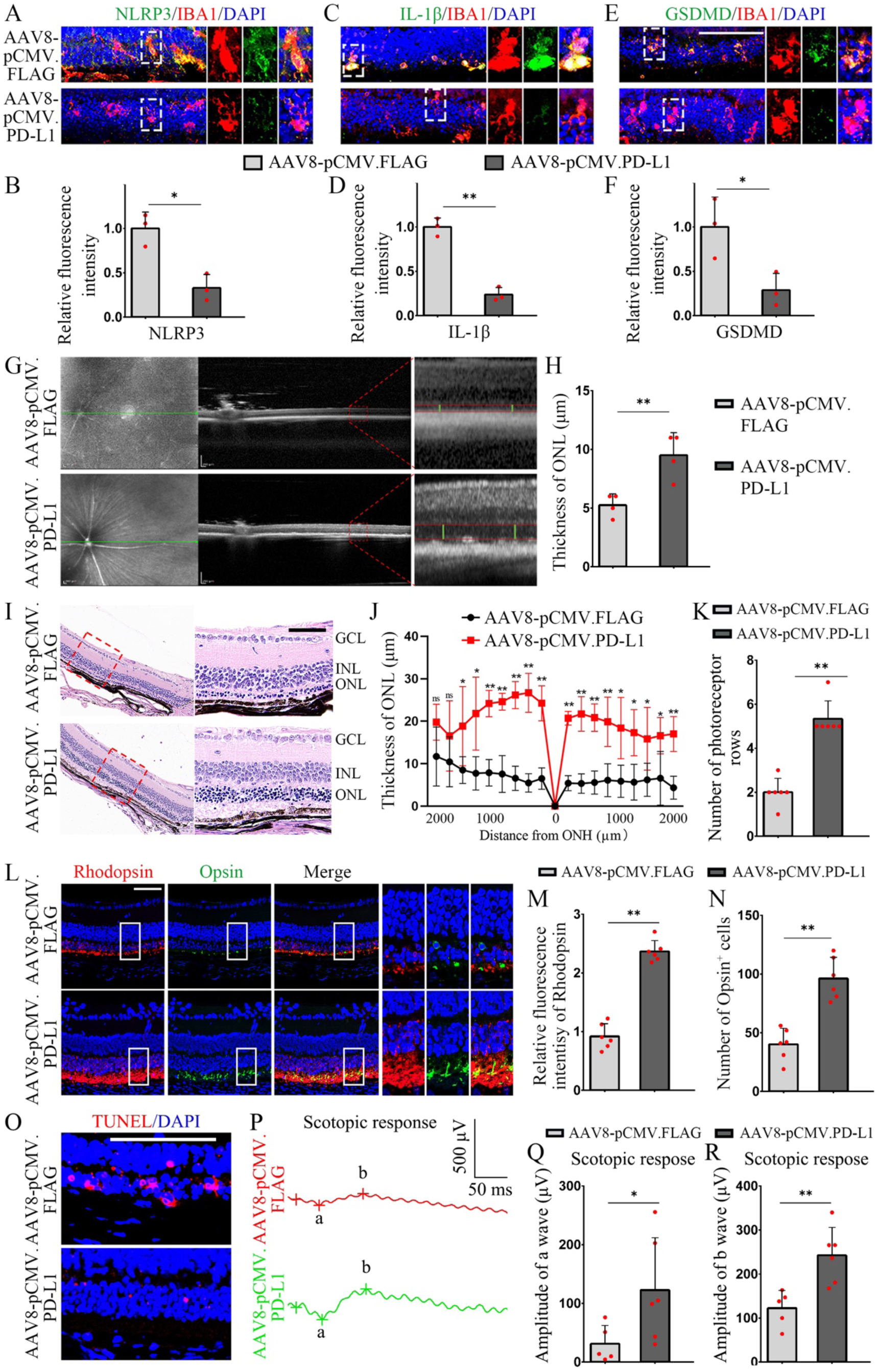
PD-L1 overexpression inhibits NLRP3 inflammasome activation and mitigates retinal degeneration in rd10 mice. All rd10 mice used in these experiments were maintained in complete darkness to slow the progression of photoreceptor degeneration. (A) Representative immunofluorescence images of NLRP3 (green) and IBA1 (red) show the reduced NLRP3 expression in microglia of AAV8-pCMV.PD-L1-infected rd10 mice at P28. (B) Quantification of relative fluorescence intensities of NLRP3 based on the results of A, n = 3. (C) Representative immunofluorescence images of IL-1β (green) and IBA1 (red) showing reduced IL-1β expression in microglia of AAV8-pCMV.PD-L1-infected rd10 mice at P28. (D) Quantification of relative fluorescence intensities of IL-1β based on the results of C, n = 3. (E) Representative immunofluorescence images of GSDMD (green) and IBA1 (red) showing reduced GSDMD expression in microglia of AAV8-pCMV-PD-L1-infected rd10 mice at P28. (F) Quantification of relative fluorescence intensities of GSDMD based on the results of E, n = 3. (G) OCT analysis showing a thicker ONL in AAV8-pCMV-PD-L1-infected rd10 mice at P32. (H) Quantification of ONL thickness from G, n = 4. (I) Representative images of H&E staining showing better preservation of ONL in AAV8-pCMV-PD-L1-infected rd10 mice at P32. (J and K) The curve diagram illustrates the thickness of ONL (J) and quantification of the number of photoreceptor rows (K) based on the results of I, n = 6. (L) Representative immunofluorescence images of Rhodopsin (red) and Opsin (green) showing increased expression and better preservation of photoreceptors in AAV8-pCMV.PD-L1-infected rd10 mice at P32. Note that the three pictures on the right were enlarged from each square of the three pictures on the left in the left-to-right order (Rhopopsin, Opsin, merge). (M and N) Quantification of Rhodopsin intensity and number of Opsin-positive cells from L, n=6. (O) Representative immunofluorescence images of the TUNEL assay show decreased apoptosis in the ONL of AAV8-pCMV.PD-L1-infected rd10 mice. (P) Representative ERG analyses show improved scotopic a-wave and b-wave amplitudes in AAV8-pCMV.PD-L1-infected rd10 mice. (Q and R) Quantifications of scotopic a-wave and b-wave amplitudes based on the results of P. AAV8-pCMV.FLAG-infected rd10 mice, n=5; AAV8-pCMV.PD-L1-infected rd10 mice, n=6. GCL, ganglion cell layer; INL, inner nuclear layer; ONH, optic nerve head; ONL, outer nuclear layer. * or ** indicates P<0.05 or P<0.01. Bar = 100 μm.

To further investigate whether PD-L1’s inhibitory effect on excessive inflammation in microglia can mitigate retinal degeneration, we used optical coherence tomography (OCT) to examine the retinal status at 18 days post-AAV infection. As shown in Figures 7G and H, the ONL was approximately twice as thick in the AAV8-pCMV.PD-L1 group compared to controls. H&E staining of retinas further confirmed that the thickness of ONL in both superior and inferior retinal regions was significantly thickened in the AAV8-pCMV.PD-L1 group compared to that of controls (Figure 7I, J), with more preserved photoreceptor rows (Figure 7I, K). Immunofluorescence also revealed that expression of Rhodopsin and Opsin in the AAV8-pCMV.PD-L1 group was significantly higher (Figure 7L, M). Additionally, while Opsin expression in control retinas appeared sparse and punctate, the AAV8-pCMV.PD-L1 group not only showed higher Opsin expression but also maintained a more normal cylindrical-shaped morphology of cone cells, resembling undamaged cells and indicating a significant preservation of retinal structure (Figure 7L, N). TUNEL assays further confirmed significantly reduced numbers of apoptotic cells in the outer nuclear layer (Figure 7O). ERG recordings of visual function showed that in AAV8-pCMV.FLAG-infected control mice, the mean amplitude of scotopic a-wave was 31.1 μV (range, 4.6-76.2 μV), and the mean b-wave amplitude was 122.5 μV (range, 63.9-163.4 μV). In contrast, the AAV8-pCMV.PD-L1-infected group exhibited significantly increased amplitudes for both scotopic a-wave and b-wave, with mean a-wave amplitude of 99 μV (range, 30.3-255.7 μV) and mean b-wave amplitude of 180.7 μV (range, 167.2-339.8 μV) (Figure 3P, Q, and R). Taken together, these data show that PD-L1 overexpression in photoreceptor cells can effectively inhibit degeneration of rod and cone photoreceptors, thereby ameliorating visual impairment.

## Discussion

It is widely accepted that microglia-driven neuroinflammation plays a key role in the progression of retinal neurodegeneration. Specialized therapeutic approaches targeting microglial reprogramming have shown promising outcomes in mitigating retinal degeneration (Scholz et al., 2015; Wang et al, 2020; Zhang et al., 2025). However, the functional diversity of microglia induced by pathogenic stimuli and their spatial distribution within retinal layers calls for the identification of broadly effective reprogramming regulators. In the present study, using a NaIO3-induced retinal degeneration mouse model, we reveal that subretinal microglia exhibit neuroprotective properties. Notably, NaIO3-stressed RPE cells upregulate the endogenous expression of PD-L1, and the overexpression of PD-L1 in RPE cells effectively reprograms microglia, attenuates NLRP3 inflammasome-mediated inflammatory responses, and slows the progression of retinal degeneration (as shown schematically in Figure 8). PD-L1 overexpression in photoreceptor cells of rd10 mice also induces a similar protective microglial reprogramming, thereby preserving photoreceptors. Altogether, these findings underscore that PD-L1 protects against retinal degeneration by suppressing inflammatory reprogramming of subretinal microglia in mouse models of experimental AMD and hereditary retinitis pigmentosa.

**Figure 8.**
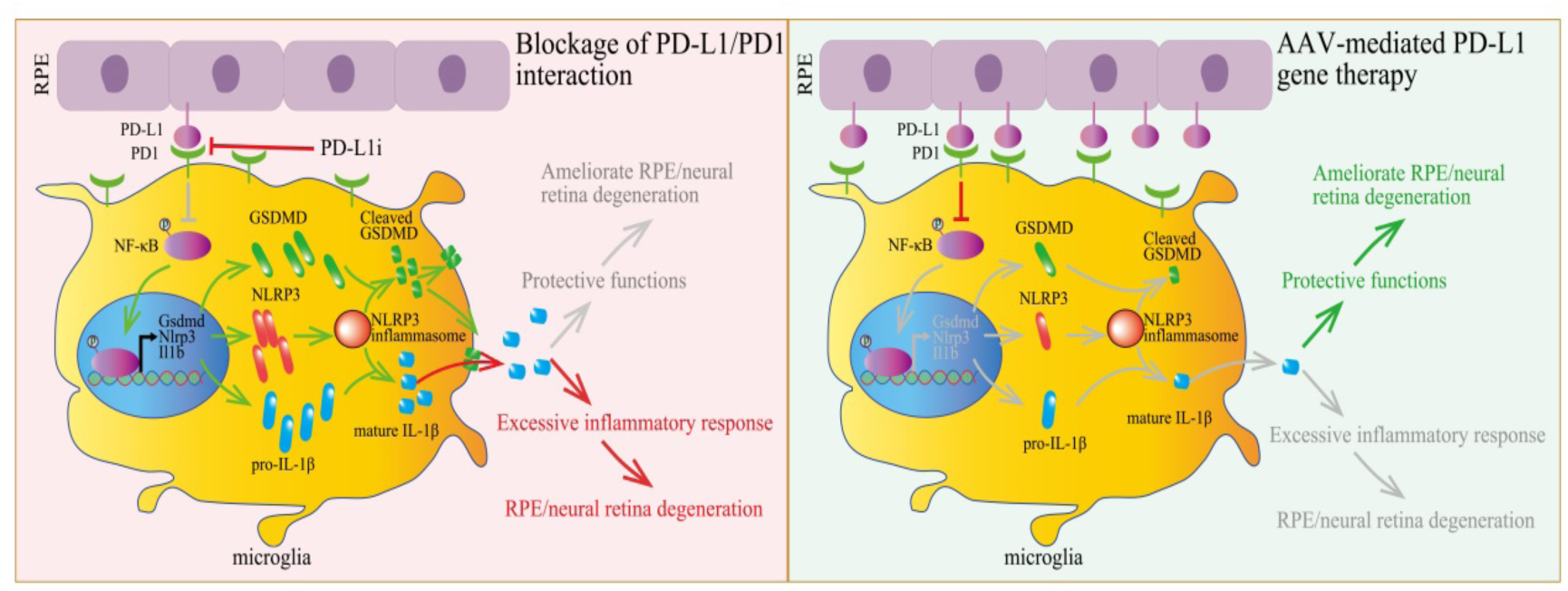
Schematic models of the effects of PD-L1/PD1 inhibition or PD-L1 gene therapy on NF-kB-mediated NLRP3 inflammasome activation. The left model illustrates that blockage of the PD-L1 and PD1 interaction enhances NF-kB activity, upregulates Nlrp3, Il1β, and Gsdmd, and leads to NLRP3 inflammasome overactivation and exacerbated retinal degeneration. In contrast, shown on the right, PD-L1 overexpression in RPE cells inhibits NLRP3 inflammasome activity and mitigates retinal degeneration.

Given these findings, PD-L1-mediated inhibition of the NLRP3 inflammasome may be a promising translational target for protecting RPE cells and photoreceptors during the progression of retinal degeneration. The pathogenic factors of retinal degeneration are complex, including congenital genetic mutations, smoking, diabetes, oxidative damage, and aging (Beatty et al., 2000; Maguire et al., 2008; Ting et al., 2017; Tomany et al., 2004). Due to this remarkable heterogeneity in etiological factors, the involved pathogenic mechanisms and regulatory networks are equally complex, encompassing oxidative stress, complement activation, lipid metabolism dysregulation, endoplasmic reticulum stress, mitochondrial dysfunction, and, notably, inflammation (Kunchithapautham et al., 2014; Fleckenstein et al., 2021). Therefore, therapeutic interventions targeting one specific form of retinal degeneration may prove ineffective in conditions driven by other pathogenic mechanisms. For instance, wet AMD therapy by anti-VEGF may be effective in reducing choroidal neovascularization and its associated leakage, thereby slowing the progression of retinal degeneration (Rofagha et al., 2013). However, this approach is inapplicable to dry AMD, accounting for 85-90% of all cases of AMD (Heier et al., 2021) and has raised clinical concerns about potentially exacerbating the progression of atrophic AMD (Enslow et al., 2016; Foss et al., 2022; Gemenetzi et al., 2017; Grunwald et al., 2014). Therefore, targeting common pathogenic mechanisms across different retinal degenerations may provide a more universally applicable therapeutic strategy. Importantly, substantial evidence suggests that microglia-mediated excessive inflammatory responses represent a critical shared pathogenic mechanism (Guillonneau et al., 2017; Krady et al., 2005; Rao et al., 2003; Tezel et al., 2022; Zabel et al., 2016). We found that AAV-mediated PD-L1 overexpression not only mitigates retinal degeneration in experimental AMD by suppressing microglial hyperactivation but also exerts similar protective effects in rd10 mice with inherited retinal degeneration. Notably, excessive microglial activation also contributes to the progression of diabetic retinopathy and experimental autoimmune uveitis (EAU) (Kinuthia et al., 2020; Okunuki et al., 2019). These findings suggest that targeting this common pathway holds promise for developing broadly effective interventions for different retinal degenerations. PD-L1-mediated microglial reprogramming may lead to a potentially versatile therapeutic strategy for preserving and restoring vision, applicable to multiple retinal degenerations, regardless of whether they are of genetic or environmental origin.

Interestingly, some studies have indicated that microglia in degenerative sites can have both beneficial and/or deleterious effects in neurodegenerative diseases. For instance, in Alzheimer’s disease, microglia can suppress disease progression through TREM2-mediated phagocytosis and clearance of Aβ and cellular debris (Hansen et al., 2018; Takahashi et al., 2005; Yeh et al., 2016). In contrast, hyperactivated microglia exhibit detrimental effects by secreting toxic factors that damage neurons and exacerbate tau pathology (Leyns and Holtzman, 2017; Liddelow et al., 2017). In cases of retinal degenerative disease, while TGF-β administration can protect against degeneration in rd10 mice (Wang et al., 2020), inhibition of the TGF-β pathway reduces choroidal neovascularization in exudative AMD models (Recalde et al., 2011; Wang et al., 2017). This discrepancy may reflect context-dependent efficacy and the wide distribution of TGF-β receptors beyond microglia, including in RPE cells (Obata et al., 1999). In addition, TGF-β promotes an epithelial-mesenchymal transition of RPE cells (Wei et al., 2022; Ma et al., 2023), which may limit its therapeutic application in diseases associated with fibrosis or neovascularization. The current study reveals a more targeted approach in NaIO3-induced retinal degeneration, where phosphorylated PD1 is localized explicitly to subretinal microglia, effectively inhibiting NF-κB signaling and NLRP3 inflammasome activation. This microglia-specific expression pattern makes the PD-L1/PD1 immune checkpoint pathway particularly attractive for precise immune modulation, providing reassurance about the potential of immune modulation in neurodegenerative diseases.

Moreover, the PD-L1/PD1 immune checkpoint generally functions to restrain T cell activation, proliferation, and cytokine production (Sun et al., 2018). Clinically, blocking this interaction enhances antitumor immunity (Brahmer et al., 2012). Our work reveals an additional role in retinal neurodegeneration: beyond regulating adaptive immunity, PD-L1 directly modulates microglial activity in the retina. Given that microglial activation facilitates leukocyte infiltration in experimental autoimmune uveoretinitis (Okunuki et al., 2019; Shirahama et al., 2025), RPE-derived PD-L1 may represent an endogenous mechanism that simultaneously counters microglial hyperactivation and adaptive immune responses. This action makes PD-L1 modulation particularly promising for slowly progressive conditions, such as age-related macular degeneration. Microglia exhibit remarkable plasticity in response to various stimuli, enabling them to play diverse roles in both health and degenerative conditions. Recent findings from single-cell sequencing demonstrate that retinal microglia comprise functionally distinct subpopulations (O’Koren et al., 2019; Yu et al., 2024), which may account for their remarkable plasticity. This heterogeneity may explain why specific interventions, such as TGF-β administration, exhibit variable effectiveness across different degeneration models. Anatomically separated subsets, including IL-34-independent microglia in the outer plexiform layer and IL-34-dependent populations in the inner nuclear layer, contribute differentially to retinal physiology (O’Koren et al., 2019). Intriguingly, both subsets can migrate to sites of injury and adapt a protective phenotype in the subretinal space (O’Koren et al., 2019; Yu et al., 2024), suggesting that RPE-derived signals can influence their inherent programming, particularly in the context of retinal degeneration. Our findings demonstrate that PD-L1 upregulation in stressed RPE reprograms these recruited subretinal microglia, while the overexpression of PD-L1 in photoreceptor cells similarly modulates microglia in the ONL of rd10 mice. This consistent activity across microglial subsets underscores PD-L1 as a potential therapeutic target with broad-spectrum efficacy.

Importantly, our study also reveals that inhibiting NF-κB transcriptional activity is pivotal for reprogramming microglia toward protective phenotypes, supported by three lines of experimental evidence: (1) Transcriptomic analyses identify NF-κB as a potential master regulator of subretinal microglial gene signatures (Yu et al., 2024) (Figure 4D, E). (2) PD-L1/PD1 signaling suppresses NF-κB activation in microglia. (3) Under PD-L1 functional inhibition, the CAPE attenuates NLRP3 priming and partially rescues degenerative pathology. NF-κB’s reprogramming effect on inflammation plays a critical role in the functional switching of various cell types. Upon activation by pathogen-associated stimuli, non-oscillatory NF-κB enhances chromatin accessibility by disrupting nucleosome protein-DNA interactions, thereby promoting the expression of genes involved in the immune response (Cheng et al., 2021). In contrast, when NF-κB activation is prevented in macrophages, an anti-inflammatory program is induced (Marwick et al., 2018). Notably, suppressing NF-κB activation enables Müller glia reprogramming into proliferative progenitors (Palazzo et al., 2020), which serve as crucial sources for retinal regeneration after injury (Goldman, 2014). Furthermore, substantial evidence confirms that NF-κB integrates multiple stress signals in neurodegeneration from oxidative damage (Broz and Dixit, 2016) to amyloid-β toxicity (Man and Kanneganti, 2015). Its activation drives both pro-inflammatory gene expression (Hikage et al., 2021; Du et al., 2022) and degenerative progression (Gao et al., 2018). Therefore, our findings highlight the potential therapeutic value of targeting NF-κB-regulated inflammatory reprogramming in microglia to prevent retinal degeneration.

Finally, NLRP3 serves as a key downstream effector and transcriptional target of NF-κB signaling. NLRP3 inflammasome complexes, well known for their role in pathogen defense, also mediate retinal stress responses. These multimolecular platforms, comprising sensors, adaptors, and inflammatory caspases, process cytokines like IL-1β and IL-18 into their active forms (Broderick et al., 2015). NLRP3 inflammasome activation in RPE cells triggers dysfunction and pyroptotic cell death (Kim et al., 2014; Gao et al., 2018), while it promotes degeneration of photoreceptors and ganglion cells (Chu et al., 2021; Viringipurampeer et al., 2016; Zhang et al., 2022). Therapeutic NLRP3 inhibition shows promise in AMD and other inflammatory retinopathies (Marneros, 2013; Kerur et al., 2013; Ambati et al., 2021). Our work positions PD-L1/PD1 as an upstream regulator of this pathway, suppressing NF-κB-dependent NLRP3 expression and subsequent inflammasome assembly in microglia. Notably, microglia-derived IL-1β stimulates RPE chemokine production (Natoli et al., 2017). However, whether the RPE-derived PD-L1 upregulation may generate a feedforward inflammatory loop to help break this production in microglia remains to be explored.

In summary, this study provides strong evidence that PD-L1 protects RPE and photoreceptor cells against NaIO3-induced retinal degeneration, as well as photoreceptors in a preclinical mouse model of retinitis pigmentosa. Our findings establish that RPE-derived PD-L1 plays a crucial role in protecting against retinal degeneration and is a potent regulator of microglial reprogramming through the regulation of the NF-κB/NLRP3 inflammasome axis. The demonstrated efficacy across multiple degeneration models supports PD-L1-targeted interventions as a potentially viable and broad strategy for preserving retinal function. Hence, this study significantly advances our understanding of the complex interactions between RPE cells and microglia in retinal homeostasis, as well as in retinal degenerative and inflammatory diseases. This approach would be even more effective when combined with other beneficial strategies, opening new avenues for future therapeutic interventions.

## Materials and Methods

### Animals

C57BL/6J mice (hereafter referred to as WT) were obtained from Beijing Vital River Laboratory Animal Technology, China. NLR family pyrin domain containing three gene knockout mice (hereafter called *Nlrp3-/-*) and rd10 mice (*Pde6b^rd10^* on the C57BL/6J genetic background) were obtained from the Jackson Laboratory. All animal experiments were carried out in accordance with the approved guidelines of the Wenzhou Medical University Institutional Animal Care and Use Committee (Permit Number: wydw 2021-0079).

For PD-L1 overexpression in rd10 mice, the mice were maintained in complete darkness to delay retinal degeneration progression, as previously reported (Cronin et al., 2012).

To induce RPE injury and retinal degeneration, mice were administered NaIO3 via tail vein injection at a dose of 20 mg/kg. At 1-, 7-, or 14-day post-injury, mice were sacrificed for eyeball sample collection.

For microglial depletion, WT mice were treated with chow mixed with 290 mg/kg of Pexidartinib (PLX3397, purchased from Selleck Chemicals, USA). After 3 days of PLX3397 feeding, retinal degeneration was induced with NaIO3. On day 7 post-NaIO3 injection (day 10 of PLX3397 feeding), mice were switched to regular chow for 7 days. Mice were treated with normal chow throughout the experiment and served as controls, and underwent the same retinal degeneration induction protocol as described above for PLX3397-treated mice.

To inhibit the interaction between PD-L1 of the RPE and PD1 of microglia, the competitive inhibitor PD-L1/PD1 inhibitor 3 (PD-L1i, purchased from Selleck Chemicals, USA) was used. PD-L1i inhibits PD-L1/PD1 binding as previously reported (Abbas et al., 2019; Du et al., 2021). Following NaIO3-induced retinal degeneration, 1 μL of 20 mM PD-L1i, dissolved in 1% DMSO in saline, was administered via subretinal injection. The control group received vehicle solution alone.

To inhibit NF-κB activity in vivo, caffeic acid phenethyl ester (CAPE, purchased from Selleck Chemicals, USA), an inhibitor of p65 nuclear translocation (Natarajan et al., 1996), was dissolved in DMSO to prepare a 150 mM stock solution, which was then diluted with saline to 1.5 mM and mixed with 20 mM PD-L1i. The control group received PD-L1i alone. One μL of the final solution was injected into the subretinal space.

### Subretinal injection

Subretinal injections were carried out as previously described (Huang et al., 2022). Briefly, the surgical procedure begins with performing a sclerotomy 1 mm posterior to the limbus using a 30-G sterilized needle. A 5 μl Ito syringe fitted with a blunt 37G Ito needle is then introduced at a 60° angle to avoid lens contact, advancing under direct visualization until retinal contact is made approximately 2-3 optic disc diameters from the optic disc. Gentle plunger pressure creates a retinotomy for subretinal access, followed by slow intermittent subretinal injection. Post-injection, fundus examination after 2-3 minutes verifies the absence of bullous retinal detachment and assesses procedural success. Finally, during anesthetic recovery on a temperature-controlled pad, antibiotic ointment is carefully applied to the cornea to prevent infection while avoiding ocular pressure or corneal damage.

### Cell culture

The human microglial HMC3 cell line was obtained from the National Collection of Authenticated Cell Cultures, China, and was cultured in MEM medium (Gibco) supplemented with 10% fetal bovine serum (Gibco) and 1% non-essential amino acids (NEAA, Gibco) in a 5% CO2 incubator. Cells were passaged every 3 days using 0.05% trypsin-EDTA.

To examine the effects of PD-L1 on HMC3 cells, recombinant human PD-L1 (Sigma-Aldrich, USA) was coated onto 6-well plates. Briefly, 1 mL of PBS containing 10 μg/mL recombinant human PD-L1 was incubated in 6-well plates at room temperature for 2 hours. The control group consisted of plates not coated with recombinant human PD-L1. HMC3 cells were cultured in PD-L1-coated 6-well plates for 48 hours before sample collection for western blot analysis.

### Analysis of gene expression profiling data

To identify protective genes potentially upregulated by RPE cells under injury conditions that may influence retinal degeneration, we analyzed previously published gene expression profiles of NaIO3-treated human RPE cell line, ARPE-19 (GSE142591, Tang et al., 2021). The online tool GEO2R was used to identify differentially expressed genes (DEGs) with at least 2-fold changes. Pathway enrichment analysis was performed using the online tool DAVID (Database for Annotation, Visualization, and Integrated Discovery) (Huang et al., 2009; Sherman et al., 2022) with the KEGG database.

To analyze highly expressed gene sets in subretinal microglia, we utilized the top 200 highly expressed genes in subretinal microglia in AMD identified through single-cell sequencing analysis as previously reported (Yu et al., 2024). Pathway enrichment analyses were performed using the online tool DAVID with the KEGG database. To further identify transcription factors that play key regulatory roles in subretinal microglial reprogramming, enrichment analysis of regulatory relationships was performed on the aforementioned gene sets using the TRRUST (Transcriptional Regulatory Relationships Unraveled by Sentence-based Text-mining) database (Han et al., 2018).

### AAV vector construction and virus injection

To achieve PD-L1 overexpression in RPE cells and photoreceptors, respectively, we engineered two recombinant adeno-associated virus (AAV) serotype 8 vectors under distinct promoter controls: RPE-specific expression system, the AAV8-pRPE65.PD-L1-GFP construct (hereafter AAV8-pRPE65.PD-L1) was generated using the GV590 backbone (GeneChem, China). The mouse Cd274 coding sequence (encoding PD-L1) was amplified via reverse transcription-PCR with primers Cd274-F1 and Cd274-R1 (Supplementary Table 1). The resulting NcoI-digested fragment was cloned downstream of the RPE-specific RPE65 promoter. AAV8-pRPE65.GFP served as the promoter-matched control vector. For pan-photoreceptor expression, we constructed AAV8-pCMV.PD-L1-FLAG (hereafter AAV8-pCMV.PD-L1) using the GV411 vector (GeneChem, China). The Cd274 coding sequence was amplified with primers Cd274-F2 and Cd274-R2 (Supplementary Table 1), followed by cloning into BamHI/NheI sites under the constitutive CMV promoter. AAV8-pCMV.FLAG was generated as the corresponding control.

All viral vectors (AAV8-pRPE65.PD-L1, AAV8-pCMV.PD-L1, and their respective controls) were packaged and purified by GeneChem (China) using standard AAV8 production protocols. For virus injections, briefly, 1 μl of AAV8-PD-L1 or the AAV8 empty virus supernatant (10^13^ genome copies/ml) was injected into the subretinal space in mice.

### Immunohistochemistry and apoptosis analysis

Immunostaining was performed following established protocols (Su et al., 2020). Briefly, cryosections were fixed in 4% paraformaldehyde (PFA), permeabilized with 0.4% Triton X-100, and blocked with 1% bovine serum albumin (BSA). Primary antibody incubation was conducted overnight at 4°C, followed by detection with either FITC- or Cy3-conjugated secondary antibodies. Imaging was performed using a Zeiss LSM880 confocal microscope. Antibody specifications are provided in Supplemental Table S2.

Apoptotic cells were identified via terminal deoxynucleotidyl transferase dUTP nick-end labeling (TUNEL) assay using the BrightRed Apoptosis Detection Kit (Vazyme, China), strictly adhering to the manufacturer’s protocol.

### Western blots and subcellular fractionation

Cytosolic and nuclear fractions were isolated as previously described (Zou et al., 2022). Retinal tissues from two mice per sample were homogenized in 500 μL ice-cold buffer I (0.3 M sucrose, 2% Tween 20, 10 mM HEPES [pH 7.9], 10 mM KCl, 1.5 mM MgCl2, 0.1 mM EDTA) and layered over 500 μL buffer II (1.5 M sucrose, 10 mM HEPES [pH 7.9], 10 mM KCl, 1.5 mM MgCl2, 0.1 mM EDTA). Fractionation was achieved by centrifugation (13,000 ×g, 4°C, 10 min), with purity verified by immunoblotting. Western blotting followed standard procedures (Wang et al., 2017), with antibodies listed in Supplemental Table S2.

### Electroretinography

Electroretinography (ERG) was recorded using a RETIport system (Roland Consult, Germany) with a custom Ganzfeld dome. Mice were dark-adapted overnight, anesthetized (ketamine/xylazine), and pupils dilated with 1% tropicamide. A corneal gold electrode recorded responses, while a tail subcutaneous electrode served as ground. Dark-adapted ERGs were elicited with flash intensities ranging from 0.01, 3 to 10 cd·s/m². Scotopic a- and b-wave amplitudes at 10 cd·s/m² were quantified for statistical comparisons. For photopic ERGs, mice were light-adapted (30 cd/m², 10 min) prior to testing to isolate cone-driven responses.

### Statistical Analysis

All experiments included ≥3 biological replicates. Data are expressed as mean ± SD. Intergroup differences were assessed using the Mann-Whitney U test, with *P* < 0.05 considered statistically significant.

## Acknowledgements

We would like to thank Dr. Heinz Arnheiter for thoughtful comments and editing of the manuscript. We also thank Drs. Tianyao Chu and Peng Zhang (Capital Medical University) for technical assistance. This work was supported by the National Natural Science Foundation of China (82101109, 8237108) and the Summit Advancement Disciplines of Zhejiang Province (Wenzhou Medical University - Pharmaceutics) and the Natural Science Foundation of Wenzhou (Y2023171).

## Author contributions

Conceived and designed the experiments: Z Su, J Wang, L Hou. Performed the experiments: Z Su, J Wang, Q Lai, Z Wang, P Liu, Z Tang, W Wang, H Yu. Analyzed the data: Z Su, J Wang, Q Lai, T Chu, P Zhang, L Hou. Contributed reagents/materials/analysis tools: Z Su, J Wang, L Hou. Funding acquisition: L Hou, J Wang, Z Su; Wrote the manuscript: Z Su, J Wang, L Hou.

