## Supplementary material for "Retinal pigment epithelium-derived PD-L1 reprograms microglial cells and protects against retinal degeneration in mouse models of experimental AMD and genetic retinitis pigmentosa": 2025 Su suppl V7.docx

**Supplementary information**

**Supplementary tables**

**Table S1. Sequences of primers used in this study.**

| Primer name | Primer sequences |
| --- | --- |
| Cd274-F1 | GGTGGATCCCGCCACGCCACCATGAGGATATTTGCTGGCATTATATTC |
| Cd274-R1 | CTCGCCCTTGCTCACCATGGTCGTCTCCTCGAATTGTGTATCATTTC |
| Cd274-F2 | GGAGGTAGTGGAATGGATCCCGCCACCATGAGGATATTTGCTGGCATTATATTC |
| Cd274-R2 | TCATCCTTGTAGTCGCTAGCCGTCTCCTCGAATTGTGTATC |

**Table S2. Antibodies used for immunostaining in this study.**

| Antibodies | Source |
| --- | --- |
| Rabbit anti-mouse IBA1 antibodies | FUJIFILM Wako Pure Chemical Corporation, Japan |
| Mouse anti-IBA1 antibodies | Abcam, UK |
| Mouse anti-Rhodopsin antibodies | Merck Limited, Germany |
| Rabbit anti-Opsin antibodies | Merck Limited, Germany |
| Anti-PD-L1 antibodies | Proteintech, USA |
| Rabbit anti-PD1 (phospho Y248) antibodies | Abcam, UK |
| Rabbit anti-phospho-NF-kB p65 (Ser536) antibodies | Cell Signaling Technology, USA |
| Mouse anti-NF-κB p65 antibodies | Cell Signaling Technology, USA |
| Rabbit anti p44/42 MAPK (Erk1/2) antibodies | Cell Signaling Technology, USA |
| Rabbit anti phospho-p44/42 MAPK (Erk1/2) antibodies | Cell Signaling Technology, USA |
| Rabbit anti pan-AKT antibodies | Cell Signaling Technology, USA |
| Rabbit anti phospho-AKT antibodies | Cell Signaling Technology, USA |
| Rat anti-NLRP3 antibodies | Thermo Fisher Scientific, USA |
| Rabbit anti-IL-1β antibodies | Abcam, UK |
| Rabbit anti-GSDMD antibodies | Abcam, UK |
| Anti-GFP antibodies | Abcam, UK |
| Anti-FLAG antibodies | Cell Signaling Technology, USA |

**Supplementary figures and figure legend**

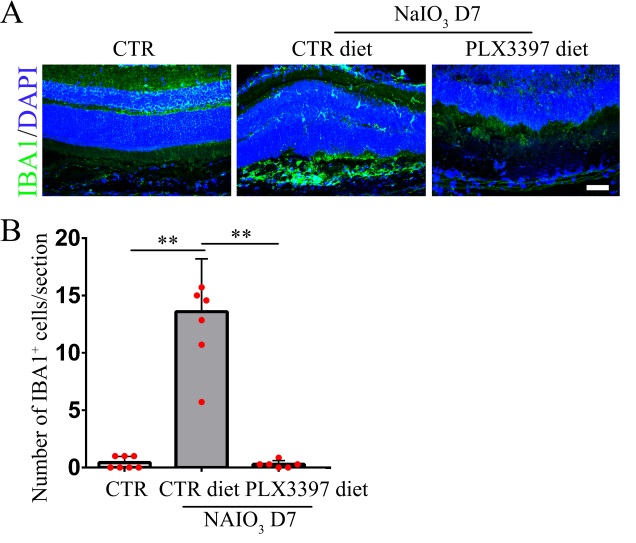

**Figure S1. Depletion of retinal microglia during retinal degeneration.** (A) IBA1 immunofluorescence staining revealed the distribution of microglia in the subretinal space. In untreated WT mice, microglia were not observed in the subretinal space. However, 7 days after treatment with NaIO_3_, many microglia were recruited into this region. Following administration of PLX3397, the CSF-1α inhibitor and the number of subretinal microglia in the NaIO_3_-injured retina were markedly reduced. (B Quantification of IBA1-positive cells based on results of A, n = 3. **P<0.01. Bar = 50 μm.

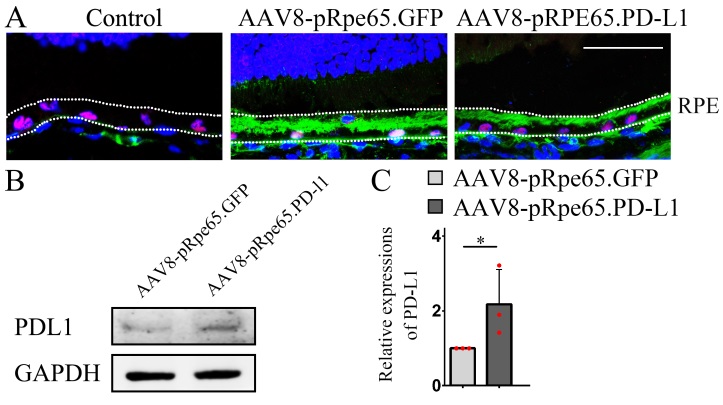

**Figure S2. AAV-mediated PD-L1 overexpression in the RPE.** (A) Representative GFP immunofluorescence images showing efficient RPE infection by AAV-pRPE65.PD-L1 and AAV-pRPE65.GFP 3 weeks post-subretinal injection. (B) Western blot analysis showing increased PD-L1 expression in AAV-pRPE65.PD-L1-infected RPE cells. (C) The bar graph shows the relative expression of PD-L1 based on the results of B, n = 3. RPE, retinal pigment epithelium. * indicates P<0.05.

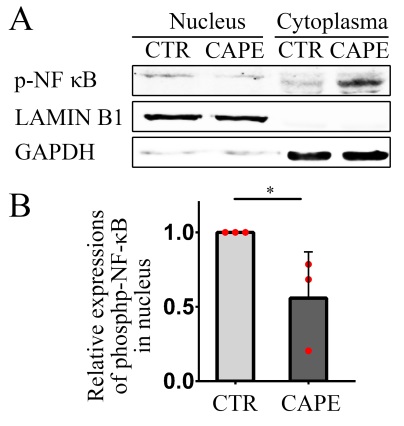

**Figure S3. Caffeic acid phenethyl ester (CAPE) inhibits the nuclear translocation of NF-κB.** (A) Analysis of subcellular localization of pNF-κB in RPE cells treated with PD-L1i by western blot analysis and (B) quantification of pNF-κB based on the results of A, n = 3. Note that CAPE leads to the inhibition of NF-κB nuclear translocation. * indicates *P*<0.05.

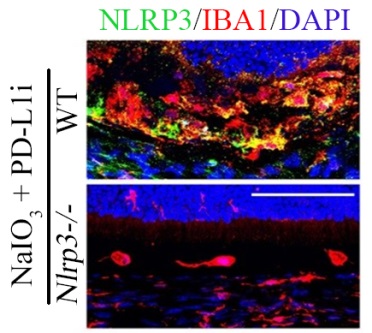

**Figure S4. Immunofluorescence analysis of NLRP3 protein in microglia**. Representative immunofluorescence images of NLRP3 in IBA1-positive microglia in WT and *Nlrp3-/-* mice. Note that there was no NLRP3 expression in *Nlrp3-/-* mice under the indicated conditions. Bar = 50 μm.

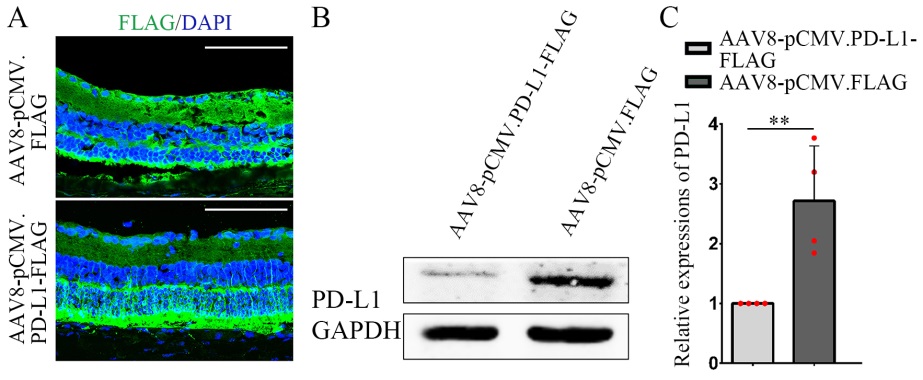

**Figure S5. AAV-mediated PD-L1 overexpression in photoreceptors.** (A) Representative immunofluorescence images of the FLAG antibody showing successful PD-L1 expression in photoreceptors 14 days post AAV8-pCMV.PD-L1 injection. (B) Representative Western blots measuring levels of PD-L1 expression in AAV8-pCMV.PD-L1-infected retinas. (C) The bar graph shows the relative expression of PD-L1 based on the results of B, n = 3. * *indicates *P*<0.01. Bar = 50 μm.
